# Computational Modeling Of Immersed Non-spherical Bodies In Viscous Flows To Study Embolus Hemodynamics Interactions For Large Vessel Occlusion Stroke

**DOI:** 10.1101/2025.03.07.642112

**Authors:** Chayut Teeraratkul, Adarsh Krishnamurthy, Debanjan Mukherjee

**Affiliations:** Paul M Rady Department of Mechanical Engineering, University of Colorado Boulder, Boulder, Colorado, USA; Department of Mechanical Engineering, Iowa State University, Ames, Iowa, USA

**Keywords:** embolisms, large vessel occlusion, stroke, fluid-particle interaction, finite elements

## Abstract

Interactions of particles with unsteady non-linear viscous flows has widespread implications in physiological and biomedical systems. One key application where this plays a fundamental role is in the mechanism and etiology of embolic strokes. Specifically, there is a need to better understand how large occlusive emboli traverse complex vascular geometries, and block a vessel disrupting blood supply. Existing modeling approaches resort to key simplifications in terms of embolic particle shape, size, and their coupling to fluid flow. Here, we devise a novel computational model for resolving embolus-hemodynamics interactions for large non-spherical emboli approaching near occlusive regimes in anatomically real vascular segment. The formulation relies on extending an immersed finite element approach, coupled with a six degree-of-freedom particle dynamics model. The geometric complexities and their manifestation in embolus-flow and embolus-wall interactions are handles using a parametric shape representation, and projection of vessel signed distance fields on the particle boundaries. We illustrate our methodology and algorithmic details, as well as present examples of benchmark cases and convergence of our technique. Thereafter, we demonstrate a parametric study of large emboli for LVO strokes, showing that our methodology can capture the non-linear tumbling dynamics of emboli originating form their interactions with the flow and vessel walls; and resolve near-occlusive scenarios involving lubrication effects around the embolus and flow re-routing to non-occludes branches. This is a key methodological advancement in stroke modeling, as to the best of our knowledge this is the first modeling framework for LVO stroke and occlusion biofluid mechanics. Finally, even though we present our framework from the perspective of LVO strokes, the methodology as developed is broadly generalizable to two-way coupled fluid-particle interaction in unsteady viscous flows for a wide range of applications.

## 1 Introduction

An embolus is a freely floating fragmented piece of blood clot or other debris (*such as cholesterol and calcific aggregates*), that can travel to a blood vessel and block or disrupt blood supply. Embolisms comprise a significant etiology for acute ischemic strokes [1]. In presence of multiple coexisting sources of embolism, it remains difficult to disambiguate embolic stroke etiology, leading to a considerable fraction of embolic strokes to be categorized as Embolic Strokes of Undetermined Sources (ESUS) [2, 3]. Additionally, small microembolic infarcts that may otherwise show no immediate symptoms, can accumulate significant tissue damage over time. This mechanism plays a central role in cognitive impairment for vascular dementia [4–6]. Owing to the fundamental role of embolic occlusions in these severe disorders, there remains a critical need to understand how an embolus originates and travels to the site of occlusion, leading to flow disruption. Addressing this need requires understanding the complex dynamic nature of embolus-hemodynamics interactions. Unfortunately, data from standard-of-care imaging, and established *in vivo* models of stroke cannot sufficiently elucidate embolus-hemodynamics interplay and embolus trajectory from it’s source to a destination vessel. Multiple *in vitro* benchtop studies using embolus analogues through a mock circulatory loop have been conducted to focus on embolus distribution across branching vessels [7–10]. These studies have elucidated several key features of embolus transport such as: size-dependence of embolus distribution; and embolus distribution being different from flow distribution across bifurcations. Despite these advancements, recapitulation of real embolism scenarios *in vitro* continues to be a challenge; and obtaining simultaneous insights on flow, embolus shape, and size effects, quantitatively within the vasculature remains difficult.

Numerical modeling techniques have provided a key alternative to address this question. Commonly, for modeling approaches in the literature, medical imaging such as contrast enhanced CT or MR angiogram has been used to reconstruct a 3D computational model of the patient vasculature, and subsequently computational fluid dynamics (CFD) simulation of blood flow and a fluid-particle interaction simulation for embolus transport is conducted through this digitally reconstructed vasculature. This approach has been used to study embolus dynamics for scenarios involving key arterial vascular segments in the cerebral vasculature [11, 12], for studying embolus transport risks from mechanical circulatory support devices [13], and for studying embolism risks in venous circulation for application in IVC filter assessment [14]. In a series of prior works [15–19], we have developed one of the most extensive computational modeling frameworks for embolus-hemodynamics interactions. In this framework, we model the embolus source to destination mapping across the entire heart-to-brain arterial pathway; and using a Monte-Carlo type sampling approach, we have generated key insights on: (a) size dependent transport patterns of emboli; (b) cardiogenic vs aortogenic vs carotid sources of emboli and how the source manifests in their distribution; and (c) the role of the Circle of Willis anastomosis and flow routing in determining embolus transport.

Efficient modeling of dynamic, non-linear, embolus hemodynamics interactions is a key complexity underlying all of these existing approaches. Fully resolving the two-way interactions of the embolus movement and the flow disturbances around the embolus within a vessel is more accurate, but is a computationally expensive task and limits the number of parametric simulation experiments for a given scenario. A commonly employed assumption in existing modeling approach to counter this challenge is that embolus-hemodynamics interactions are one-way coupled, meaning that the flow influences the embolus trajectory, but the embolus itself does not significantly alter the flow. As demonstrated in prior works [15, 17], this is a reasonable assumption for small-to-medium embolus sizes in comparison to vessel diameters. However, at larger embolus-to-vessel diameter ratio, this assumption starts breaking down, making model predictions to be insufficient at capturing true embolus-hemodynamics interactions. This assumption is accompanied by a second geometric simplification wherein the embolus is assumed to be spherical in shape. Again, while for small-to-medium emboli, departures from spherical shape may not be as significant, same cannot be inferred for larger emboli, where interactions of an arbitrarily shaped embolus with curved and tortuous vessel walls may lead to non-intuitive embolus pathways and distribution patterns. Presently, all *in silico* embolus modeling efforts rely on studying spherical shaped embolic analogues traversing within vessels such that embolus to vessel size ratios are small-to-medium.

Consequently, there remains a need to develop modeling techniques that are suitable for scenarios involving large arbitrarily shaped embolic fragments in supplying vessels into the brain. Clinically, this is of high relevance in not only resolving biophysical phenomena and transport processes happening local to the occlusion site at the vessel of interest, but also in developing *in silico* models for a specific category of ischemic events categorized as Large Vessel Occlusions (LVOs). LVOs comprise around 30 % of ischemic stroke cases [20], but are known to cause significant neurological deficits owing to their large infract size [21], and often recognized as disproportionate contributors to stroke morbidity and mortality [22]. There presently exists no modeling approach suitable to handle LVO scenarios. Another key clinical scenario presenting critical challenges involves distal embolization during mechanical thrombectomy [23], where the risk of arbitrary shaped fragments of varying sizes embolizing within the brain is associated with reduced treatment outcomes, but remains challenging to computationally recapitulate. Motivated by these state-of-the-art needs, here we describe a novel computational model for embolus-hemodynamics interactions where: (a) we introduce an immersed domain approach for two-way embolus-hemodynamics coupling, going beyond the size limits for one-way coupled interactions; (b) use an analytical geometric shape family to represent embolus geometries and their departure from sphericity; and (c) introduce a novel methodology using signed distance fields to capture complex embolus-wall interactions in a vessel especially for arbitrary embolus shapes. We develop the methodological components, demonstrate their performance, and illustrate the overall methodology computational models of embolus movement through pulsatile flow within carotid artery bifurcation models.

## 2 Methods

### 2.1 An immersed finite element approach for embolus hemodynamics interactions

For modeling unsteady embolus-hemodynamics interactions, we adopted an immersed finite element method (IFEM) based approach, based on the formulation originally described in [24] and further modified in [25–27]. Immersed domain techniques such as IFEM are particularly advantageous for this application because they do not require the fluid mesh to conform to the embolus domain. Specifically, as the embolus travels a large distance from its source to a destination vessel, mesh conforming methods will result in highly distorted mesh elements which will necessitate computationally expensive re-meshing. Conversely, within the IFEM framework, the fluid mesh remains static, and the solid embolus domain is tracked in a Lagrangian manner. The interaction force is computed on the solid domain and projected onto the background fluid domain, with some customizations that we outline here. A detailed review of the mathematical foundations of these methods is beyond the scope of this presentation. For a comprehensive description of IFEM pertaining to deformable structures, we refer the reader to [24–27]. In our IFEM formulation, we assumed that the background fluid (*that is, blood*) occupies the entire computational domain Ω, while the solid (*embolus*) domain Ω^*s*^, occupies a finite volume within Ω. We refer to the region where Ω^*s*^ overlaps with Ω as the *“artificial”* fluid domain, denoted as 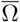 and the domain 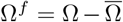 as the true fluid domain. Our numerical methodology involves solving a modified version of the fluid momentum equations within Ω, and subsequently constraining the velocity field within 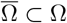 to match the velocity field in Ω^*s*^ and satisfy the no-slip boundary condition. For the formulation presented here, we assume that embolus deformations remain small, and each individual embolus domain is modeled as a rigid body with 3-dimensional rotational and translational degrees of freedom (*often referred to also as a 6 degree-of-freedom model*). Our approach here, therefore, is based on the formulation details for the semi-implicit variant of the IFEM as described in [26, 27]. For our numerical formulation, assuming ***<u>u</u>*** and *p* be the velocity and pressure field defined on the total computational domain Ω, we can state a momentum balance equation:

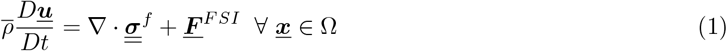

and assuming blood to be a constant density incompressible fluid, we have the mass balance relation:

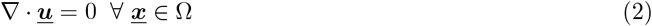

where ***<u>F</u>*** ^*F SI*^ is the fluid structure interaction force defined on Ω^*s*^. For a general case of a deformable solid, the force ***<u>F</u>*** ^*F SI*^ is defined as:

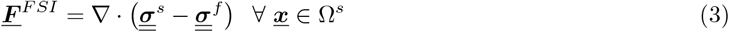

where 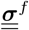 is the fluid stress and 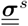 is the solid internal stress. For this application, we assumed that blood is a Newtonian viscous fluid, therefore

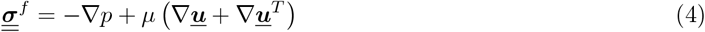

where *µ* is the bulk dynamic viscosity of blood, and *p* depicts the pressure. Since here we assume that the embolus is a rigid body, we enable two key simplifications to the original IFEM equations. First, typically, the solid internal stress 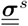 is defined based on a chosen solid constitutive law. However, under a rigid solid assumption, there is no internal stress within the solid, so 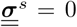. Second, in the IFEM framework described in [27], the continuity equation within 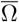 accounts for non-zero compressibility of the immersed solid. However, because our solid is rigid, we can assume the divergence-free condition for the entire domain, substantially simplifying the formulation. To distinguish the solid and fluid domains, we define ℐ (***<u>x</u>***) as a scalar indicator field for 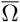 within Ω^*s*^ such that:

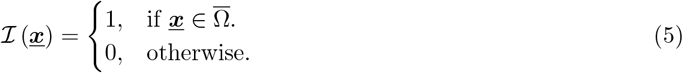

and subsequently, 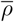 is defined as a discontinuous density field:

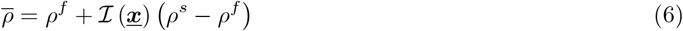

For brevity of notation, we will henceforth drop the “(***<u>x</u>***)” from ℐ. Additionally, since ℐ represents the artificial fluid region 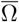 as a discontinuous scalar field, a large density ratio between the fluid and solid can lead to numerical instability, as discussed in [26]. To address this issue, we introduce a transitional region in ℐ (*a regularization approach*), which smoothly varies from 0 to 1, following the approach in [26]. This indicator field is updated at each time step based on the embolus’s position and orientation. Equations (1) and (2) are then solved using a Petrov-Galerkin stabilized finite element algorithm. Specifically, numerically we find solutions (*trial functions*) ***<u>u</u>*** ∈ 𝒮 and *p* ∈ ℒ_2_(Ω), with test functions ***<u>w</u>*** ∈ 𝒱 and *q* ∈ ℒ_2_(Ω) such that:

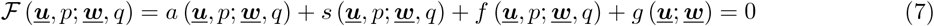

In the above: 𝒮 denotes the trial solution space, and 𝒱 denotes the test function space, such that:

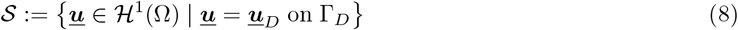

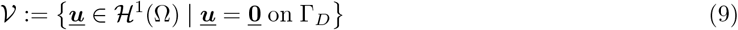

and ℒ_2_(Ω) denotes the space of square integrable functions. The first term *a*(.) in Equation (7) represents contribution from the semi-discrete Navier-Stokes formulation, detailed as follows:

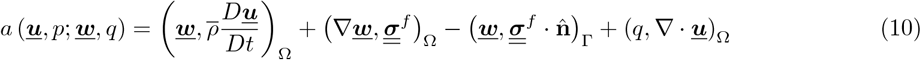

where we have used the simplifying notation (*a, b*)_Ω_ := ∫_Ω_ *a · b d*Ω. The second term in Equation (7) represents the stabilization contributions comprising the Petrov-Galerkin stabilization for the convection dominant flow regimes [28], stabilization for linear pressure-velocity interpolation space [28], and backflow stabilization commonly used in pulsatile hemodynamics simulation [29]. The contributions are detailed as follows:

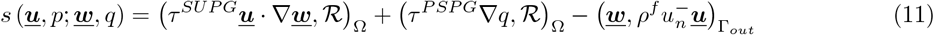

where ℛ is the residual of the momentum and mass balance equations (1) and (2). The first two terms are the contribution from the Petrov-Galerkin stabilization with the stabilization parameters *τ* ^*SUPG*^ and *τ* ^*PSPG*^ being defined following [30] as:

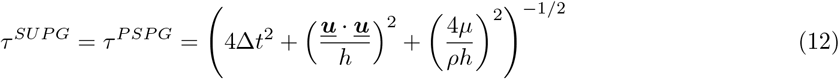

The final term in Equation (12) denotes the backflow stabilization contribution which stabilizes error buildup at the outlet due to large flow reversal. In Equation (11), 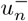 is defined as:

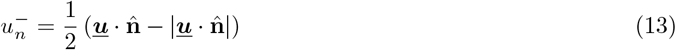

where 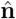 is the outward normal direction of the outlet, resulting in backflow stabilization being non-zero only when the wall normal component of the flow velocity points inward. Following prior works [29], we set the stabilization strength *β* = 0.2 for all of our simulations. The third and fourth terms in Equation 11, represent the fluid-solid interaction contributions, requiring some key customizations. First, *f* (***<u>u</u>***, *p*; ***<u>w</u>***, *q*) denotes the contribution from the fluid-structure interaction force (***<u>F</u>*** ^*FSI*^) from the immersed rigid solid:

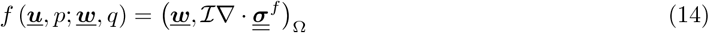

To reduce the continuity requirement for the velocity approximation space, we use integration by parts to re-formulate *f* (***<u>u</u>***, *p*; ***<u>w</u>***, *q*):

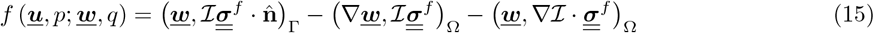

We note that regularization of the discontinuous ℐ function (*as noted earlier in the description*) enables the above integration to hold true. ℐ is only non-zero inside Ω because Ω^*s*^ is immersed within Ω. Thus, Equation (15) becomes:

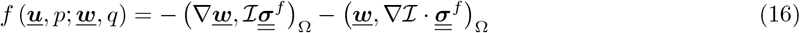

Finally, *g*(***<u>u</u>***; ***<u>w</u>***) represents numerically constraining the velocities, where we employ a fictitious domain penalization approach as outlined in [30–32]. Specifically, we state:

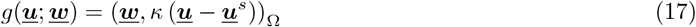

where the term *κ* represents a consistent penalty scale factor, formulated as:

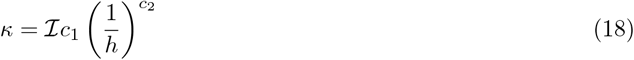

The resulting contribution from Equation (18) can effectively constrain the velocity with Ω to the velocity of the solid domain ***u***^*s*^ in a fictitious domain sense, acting nodally without the requirement of body-conformed discretizations, as previously demonstrated in [30–32].

### 2.2 Parametric modeling of arbitrary embolus shapes

A critical modeling component here is the representation of the embolus domain (Ω^*s*^) shape. Here, we choose to use an analytical family of geometries named superquadrics [33–35]. We have previously demonstrated that superquadric geometries offer substantial numerical flexibility in modeling complex shapes for biomedical and expecially vascular flows with thrombotic applications, with possibilities of constrained shape modeling, and parametreic variations in shape geometry [30]. Specifically, an individual superquadric shape can be uniquely represented using three size parameters (*a*_1_, *a*_2_, *a*_3_), and two shape or roundedness parameters (*ϵ*_1_, *ϵ*_2_). Points on a superquadric with the above parameters, in their reference configuration, are represented using an analytical function 𝒫 as follows:

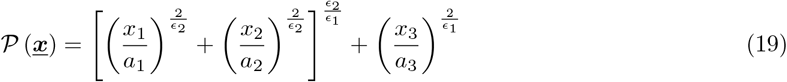

such that 𝒫 (***<u>x</u>***) *<* 1 if the point ***<u>x</u>*** is inside the superquadric solid object, 𝒫 (***<u>x</u>***) = 1 if the point ***<u>x</u>*** is on the boundary of the object, and 𝒫 (***<u>x</u>***) *>* 1 if the point ***<u>x</u>*** is outside the object. In a transformed coordinate space, that is, current configuration, the superquadric analytical shape function can be re-stated as follows:

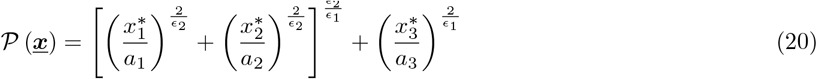

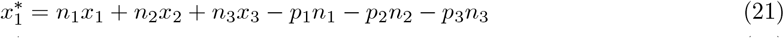

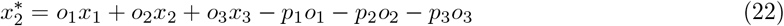

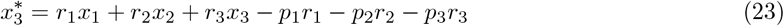

where the transformation from reference to current configuration is represented using a combination of rotations (*n*_*i*_, *o*_*i*_, *r*_*i*_) and a translational displacement *p*_*i*_ respectively as follows:

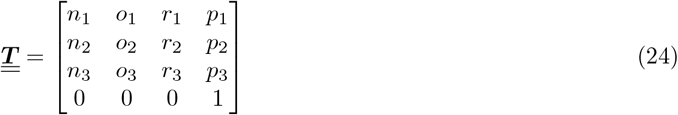

With this shape representation, we can directly write out the function 𝒫 for an embolus as function of its translational and rotational degrees of freedom, and mark element nodes or Gauss quadrature points using the indicator function defined in Equation (5). Additional mathematical details on using superquadrics for thrombotic and embolic phenomena is provided in [30]. For demonstration here, we choose families of shapes such that *ϵ*_1_ = *ϵ*_2_ = 1, leading to superellipsoid shaped emboli. However, since the IFEM formulation is linked to the shape using the function ℐ which in turn is directly linked to 𝒫, we can extend to any arbitrary shape embolus without additional complexity.

### 2.3 Embolus dynamics as a 6 DOF rigid body motion

The dynamics of each embolus, under the assumption thet they are nearly-rigid bodies, is modeled using linear and angular momentum conservation around the embolus center of mass ***<u>x</u>***_*c*_. Thereby, the 6 rotational and translational degrees of freedom in 3 dimensional dynamics of the embolus are tracked using the following equations:

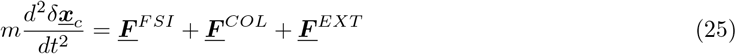

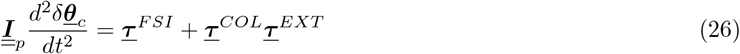

where *δ****<u>x</u>***_*c*_ is the displacement of the center of mass ***<u>x</u>***_*c*_ and *δ****<u>θ</u>***_*c*_ is the angular displacement about ***<u>x</u>***_*c*_; *m* and 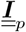 are the mass and mass moment of inertia of the solid; ***<u>F</u>*** ^*F SI*^ and ***<u>τ</u>*** ^*F SI*^ are the force and torque applied to the embolus by the fluid; ***<u>F</u>*** ^*COL*^ and ***<u>τ</u>*** ^*COL*^ are the force and torque due to wall collision, and finally ***<u>F</u>*** ^*EXT*^ and ***<u>τ</u>*** ^*F SI*^ are the externally applied force and torque applied to the embolus, e.g., gravity. The fluid forces and torque applied to the solid domain in Equations (25) and (26) is the surface integral of the traction applied by the fluid on the solid surface such that:

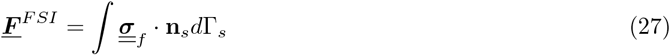

where **n**_*s*_ is the outward facing normal on the embolus boundary. Similarly, the torque due to fluid forces is computed as:

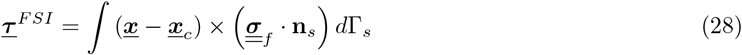

As an arbitrary shaped embolus interacts with the flow and the wall, it is necessary to correctly represent the 3-dimensional orientation of the embolus over time within the vessel. Applying rotations to a 3D solid body by sequentially rotating it around each Cartesian axis using Euler angles can lead to Gimbal lock (*a well known artefact in rigid body dynamics*), where the rotations can degenerate into two dimensions. To address this issue, in our method rigid body rotations were represented using quaternions. Quaternions are four-element complex numbers which represent three-dimensional rotations about an arbitrary axis [36, 37]. Mathematically, a single quaternion is represented as a vector of four real numbers:

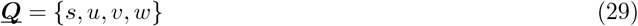

such that, a quaternion representing rotation angle *θ* about an axis ***t*** = *{t*_1_, *t*_2_, *t*_3_*}* is represented as:

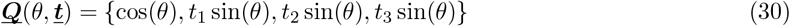

The product of two quaternions represents successive rotations, allowing a series of rigid body rotations to be represented as a single quaternion value. Specifically, here we store the dynamic orientation of an embolus as a quaternion. At each time step, angular displacement was applied by multiplying a quaternion representing the angular displacement with the current quaternion value that represents the embolus orientation. A new orientation quaternion ***<u>Q</u>***_3_ representing a series of consecutive rotations first by quaternion ***<u>Q</u>***_1_ = *{s*_1_, *u*_1_, *v*_1_, *w*_1_*}* then quaternion ***<u>Q</u>***_2_ = *{s*_2_, *u*_2_, *v*_2_, *w*_2_*}* is computed as a product of ***<u>Q</u>***_1_ and ***<u>Q</u>***_2_ as:

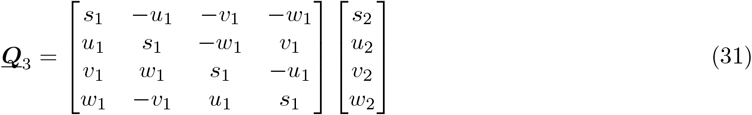

Hence, the quaternion at the *n*^*th*^ time step in the simulation was given by *Q*_*n*_ = *Q*_0_ \**Q*_1_ *… \**Q*_*n*−1_. Similar to three-dimensional rotations, quaternion multiplication is not commutative. Therefore, the order of rotations must be preserved for computations as proposed here. Finally, at each time-step the quaternion *Q* = *{s, u, v, w}* was converted to Euler angles ***<u>θ</u>*** = *{θ*_*x*_, *θ*_*y*_, *θ*_*z*_*}* using the following mathematical relations:

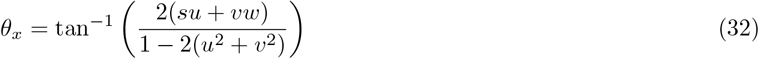

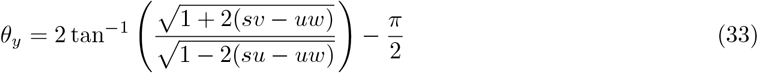

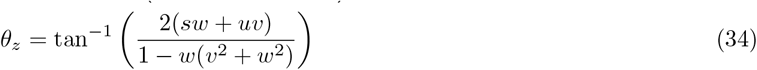

where the Euler angles rotation is applied first about the *z*− axis, then the *y*− axis, and then the *x*−axis. Once updated, the Euler angles are used to create the rotation matrix that provides the components (*n*_*i*_, *o*_*i*_, *r*_*i*_) to transform the embolus domain using the equations described in Section 2.2. In our formulation, Equations (25) and (26) are integrated in time using Gear’s algorithm which is an explicit time marching scheme commonly used in discrete element simulations [38, 39]. Gear’s algorithm consists of a single-step predictor-corrector scheme, where velocities and accelerations are first predicted for the time step *t* +Δ*t* based on their values at time *t*; and then updated using a correction based on time-step size Δ*t*. Here, we used a fifth order Gear’s scheme that is commonly employed in discrete element methods and molecular dynamics [38, 39]. Additional numerical details about the predictor-corrector steps are provided in Supplementary Material.

### 2.4 Evaluating immersed FSI force using a non body-fitted approach

Computation of ***<u>F</u>*** ^*FSI*^ and ***<u>τ</u>*** ^*FSI*^ in Equations (27) and (28) requires integration of the fluid stresses along the immersed solid boundary Γ_*s*_. An accurate resolution of the FSI force is particularly important because flow-induced forces will directly determine the trajectory of the embolus. This necessitates a mapping of the fluid stress 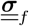 from Ω to Γ_*s*_, requiring a robust interpolation of the pressure value and the gradient of the fluid velocity onto Γ_*s*_ despite a non-conforming or non-fitted discretization of the immersed boundary. To obtain an accurate interpolation of pressure and velocity gradient comprising 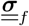 on the immersed Γ_*s*_, we use the mathematics of Reproducing Kernel (RK) interpolation - a kernel-based interpolation method [40, 41] which is frequently used in immersed finite element applications [25–27, 42]. Here, we briefly outline the RK interpolation method as implemented for our application. For a more detailed elaboration of the RK basis functions, we refer the reader to [41]. Briefly, in our approach we defined a function on the solid boundary *f* ^*h*^(***<u>x</u>***^*s*^) as the weighted sum of the function value evaluated on the fluid points ***<u>x</u>***^*f*^ as follows:

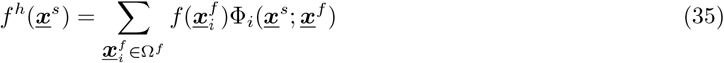

where Φ_*i*_ denotes the reproducing kernel basis function. Additionally, the gradient of *f* (***<u>x</u>***^*f*^) was interpolated onto the solid boundary as follows:

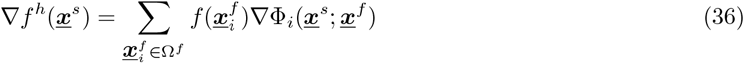

The RK basis function Φ_*i*_(***<u>x</u>***^*s*^) can be in turn defined as follows:

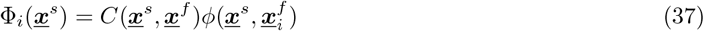

where *C*(***<u>x</u>***^*s*^, ***<u>x</u>***^*f*^) is the correction function and 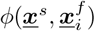 is an interpolating kernel function with compact support. In this implementation, we use the cubic spline kernel interpolation function defined as follows:

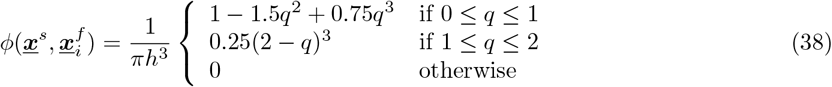

where *h* defines the support size of the cubic spline and 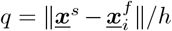 is the normalized distance measure from the solid point to the fluid point. The coefficient 1*/πh*^3^ is used so that ∫ *ϕd*Ω = 1 in 3 dimensions. Because *ϕ* has a finite support, Equation (35) is only summed over the neighboring fluid points of ***<u>x</u>***^*s*^, thereby reducing the computational cost significantly. Finally, the correction function *C*(***<u>x</u>***^*s*^, ***<u>x</u>***^*f*^) is introduced to ensure that the RK interpolator reproduces a certain polynomial degree exactly [40]. In this work, we choose a linearly complete RK function where *C* is obtained by solving the following system of linear equation:

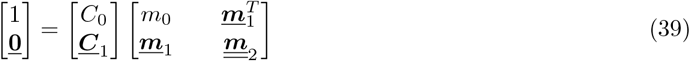

where the moments *m*_0_, ***<u>m</u>***_1_, and 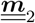 are mathematically defined as:

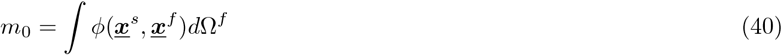

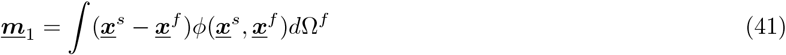

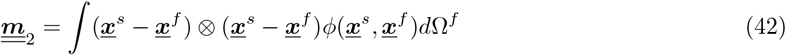

We note that for an interpolation of a vector-valued function, this interpolation procedure is done component-wise. While RK interpolation can obtain accurate gradient estimation, it is possible that the fictitious domain constraint results in a ”fattened/widened” particle or embolus boundary region due to the lack of an explicit boundary where DOF points are defined. This effect has been noted in a prior work in [30] where the viscous drag force convergence with respect to mesh size is shown to be sub-optimal. To alleviate this problem, we evaluate 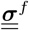 (*through RK interpolation*) on solid points one cell length away from the solid point such that:

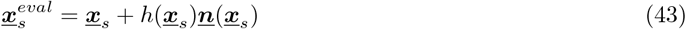

where *h*(***<u>x</u>***_*s*_) is the size of the cell which contains ***<u>x</u>***_*s*_ and ***n***(***<u>x</u>***_*s*_) is the embolus boundary normal at point ***<u>x</u>***_*s*_. This evaluation point relaxation is still consistent because as *h*(***<u>x</u>***_*s*_) approaches 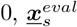 approaches ***<u>x</u>***_*s*_. Finally, while Equation 11 is solved over the entire domain Ω, ***<u>F</u>*** ^*FSI*^ is interpolated solely from points within Ω^*f*^. This is because points within 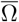 are artificial and do not correspond to real fluid point and therefore are treated as holes within the mesh. This approach is analogous to using RK interpolation to approximate forces from a simulation where the mesh explicitly conforms to the embolus boundary. In such a case, Ω does not exist, and RK interpolation is perform on points near the mesh boundary conforming to the embolus. RK interpolation is shown to retain good interpolation accuracy for both the function and the function’s normal gradient near a boundary as demonstrated in the *Boundary point* cases in Figure 3.

### 2.5 Evaluating wall interactions using signed distance field

For scenarios comprising large embolus size (*in comparison to vessel diameter*) and arbitrary non-spherical shape, robust treatment of embolus interactions with vessel walls is critical. Yet, checking and resolving contact between an arbitrary shaped embolus considering realistic patient-specific vascular geometries with significant curvature, tortuosity, and diameter variation, is a challenging task with significant computational expense. Here, we propose a novel wall interaction modeling approach using a geometric function called the Signed Distance Field (*SDF*), which calculates the orthogonal distance of a given point to a closed boundary [43, 44]. Here, we model the embolus-vessel collision as a semi-rigid collision. The collision force between the embolus and the vessel wall is modeled as a function of the overlap distance between the embolus and the wall boundary without having to solve the actual deformation on the embolus and vessel wall, hence significantly saving on computational cost. Semi-rigid collision assumption is commonly employed in discrete element simulations and is generally valid for small overlap distance [39]. For our methodology, we define the embolus-vessel collision force as:

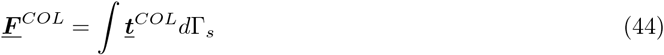

where ***<u>t</u>***^*COL*^ is a collision force per unit area being applied on the embolus wall. This is, therefore, an effective distributed surface traction applied along the embolus boundary to account for wall collision. Since our focus here was on developing the overall formulation for embolus-hemodynamics-wall interplay, we employed a simple approach equivalent to a non-linear spring, as outlined in several studies [45]. Based on this, we define:

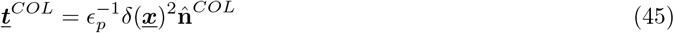

where *δ*(***<u>x</u>***) is the penetration depth at point 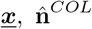 is the wall normal direction, and *ϵ*_*p*_ is a spring parameter that is chosen to be a small number 1.0e-6 to penalize any embolus-wall inter-penetration. The torque applied on to the embolus due to the wall collision is then computed as:

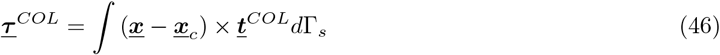

For mathematically obtaining ***<u>t</u>***^*COL*^, we compute the SDF for the vessel wall along the mesh nodes of the flow domain, and then at each time instance we project the SDF and the gradient of the SDF onto points along the embolus boundary. For analytical embolus domain representations these points can be regularly spaced as per intrinsic superquadric surface parameterization [33]. For embolus geometry represented as a triangulated surface, the SDF can also be projected directly onto the quadrature points along the triangles - which is the approach we chose for the simulations shown here. The sign in *SDF* indicates whether a point is inside or outside the enclosing boundary. In our implementation, points inside the boundary are assigned a negative value, while points outside are assigned a positive value. Finally, the normalized SDF gradient gives the normal at the wall:

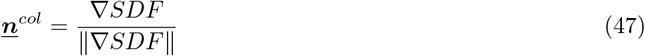

Defining, *δ* then as follows:

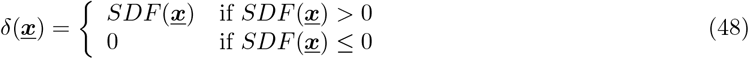

which can be plugged back into Equation (45) to get the collision interactions. The grid interpolation and projection of the SDF on the embolus boundary are computationally inexpensive, making the overall collision check numerically robust yet cheap. In this work, we use the *SDF* implementation from the open-source computational geometry library libigl [46, 47].

## 3 Numerical tests with the proposed method

### 3.1 Convergence study of the RK interpolation

We demonstrate the convergence behavior of the RK interpolation procedure of a function defined on fluid points ***<u>x</u>***_*f*_ interpolated onto a solid point ***<u>x</u>***_*s*_. The procedure is illustrated in Figure 2. To achieve this, first we construct uniformly spaced points with spacing size denoted as *h* as shown in Figure 2.a. To mimic degree-of-freedom (DOF) points typically obtained from an unstructured fluid mesh, each point within the uniform grid is randomly perturbed by a uniformly random distance between 0 and *h/*2 resulting in a point cloud illustrated in Figure 2.b. This sampling procedure is repeated for varying initial grid spacing *h* for both 2D and 3D domain geometries, and these points will subsequently serve as the background fluid points where the function values (*velocity and pressure*) are defined. Here, we examine the convergence behavior of RK interpolation on two sample points: the first point is located at ***<u>x</u>***_*center*_ = (0.5, 0.5) in 2D and ***<u>x</u>***_*center*_ = (0.5, 0.5, 0.5) in 3D representing a solid point located within the fluid domain far away from the boundary. The second point is located at ***<u>x</u>***_*boundary*_ = (0.5, 0) in 2D and ***<u>x</u>***_*boundary*_ = (0.5, 0, 0.5) in 3D representing an interpolation on a solid point close to the fluid domain boundary. Figure 3 illustrates the convergence rate of RK interpolation of the function *f* = sin(2*πy*) and its derivative showing that the function interpolation error decreases as *O*(*h*^3^). Furthermore, we examine the wall normal gradient interpolation behavior in this sample problem by computing ∂_*y*_*f* using Equation (36) showing a convergence rate of *O*(*h*^2^). This is analogous to interpolating the normal component of the strain rate term, for instance, which makes up the viscous part of the stress tensor leading to the flow-induced force. Based on these, we established that the RK interpolation procedure can accurately interpolate a function and a wall-normal gradient well in both in the middle or interior of the domain and near the fluid domain boundary.

**Figure 1:**
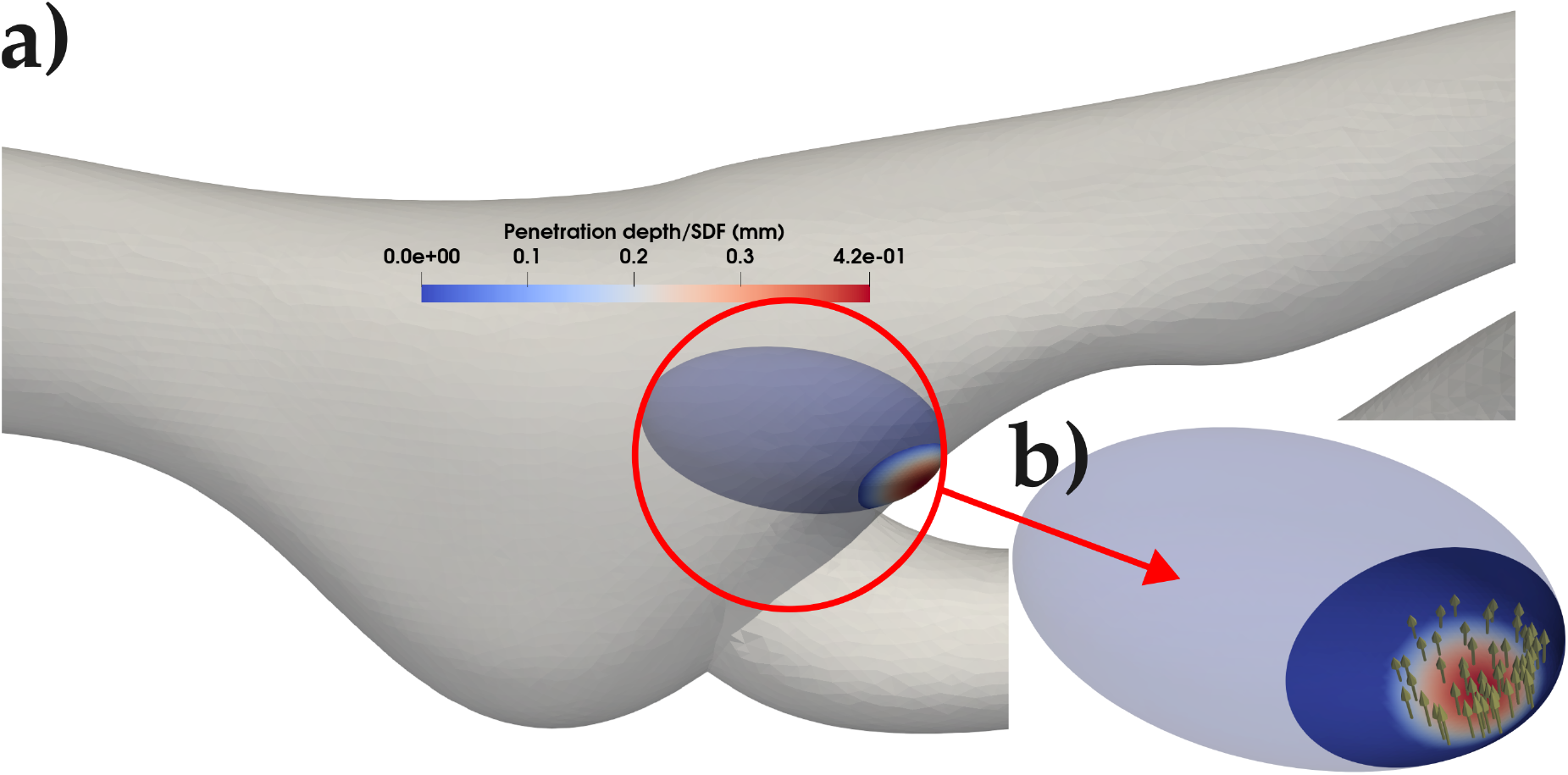
Simplified illustration of the contac resolution scheme using signed distance field of the vessel wall projected on the particle (embolus) boundary (as indicated in panel a.). The resulting contact penetration distance is used to compute a distributed interaction force along the wall of the particle/embolus (as shown in panel b.)

**Figure 2:**
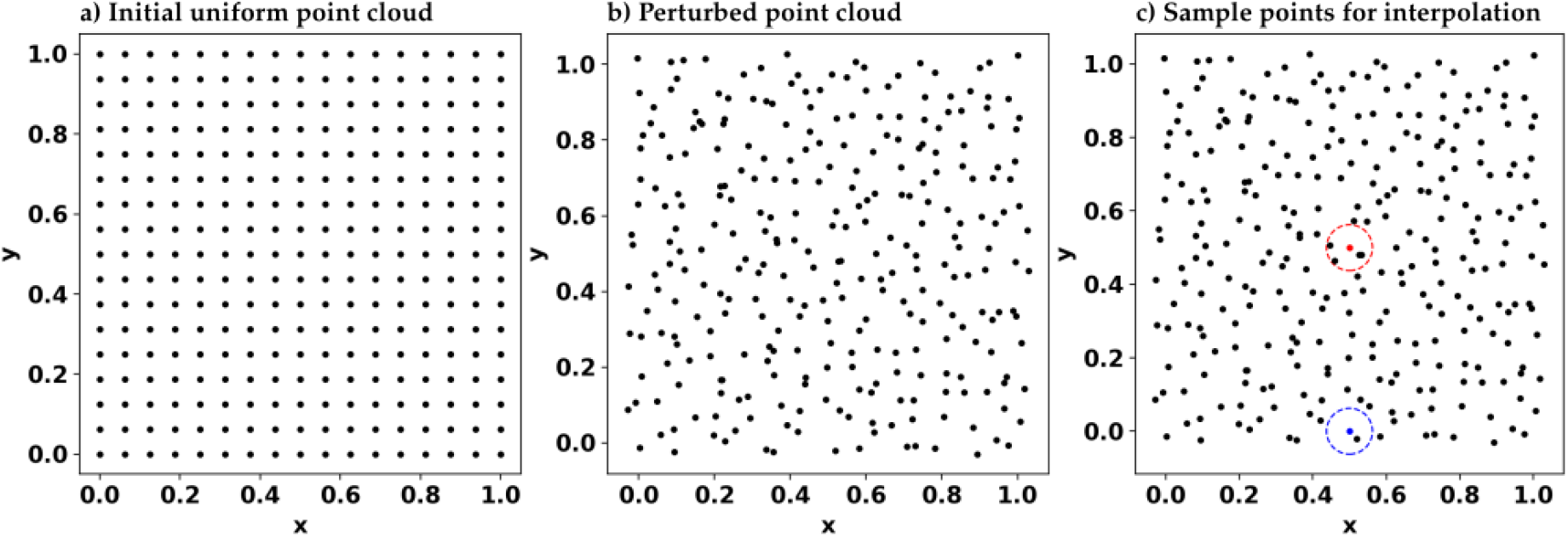
Sample workflow illustrating the RK point cloud interpolation convergence study. Panel a) the point cloud is initially seeded in an uniformly spaced Cartesian grid with spacing h; b) the uniform grid is randomly perturbed by distance ± h/2 to mimic point cloud obtained from an unstructured grid; c) interpolation convergence is studied on two sample points: first point is located at **<u>x</u>** = (0.5, 0.5) (red) representing a point far from the domain boundary, and the second point is located at **x** = (0, 0.5) (blue) representing interpolation of points near a domain boundary.

**Figure 3:**
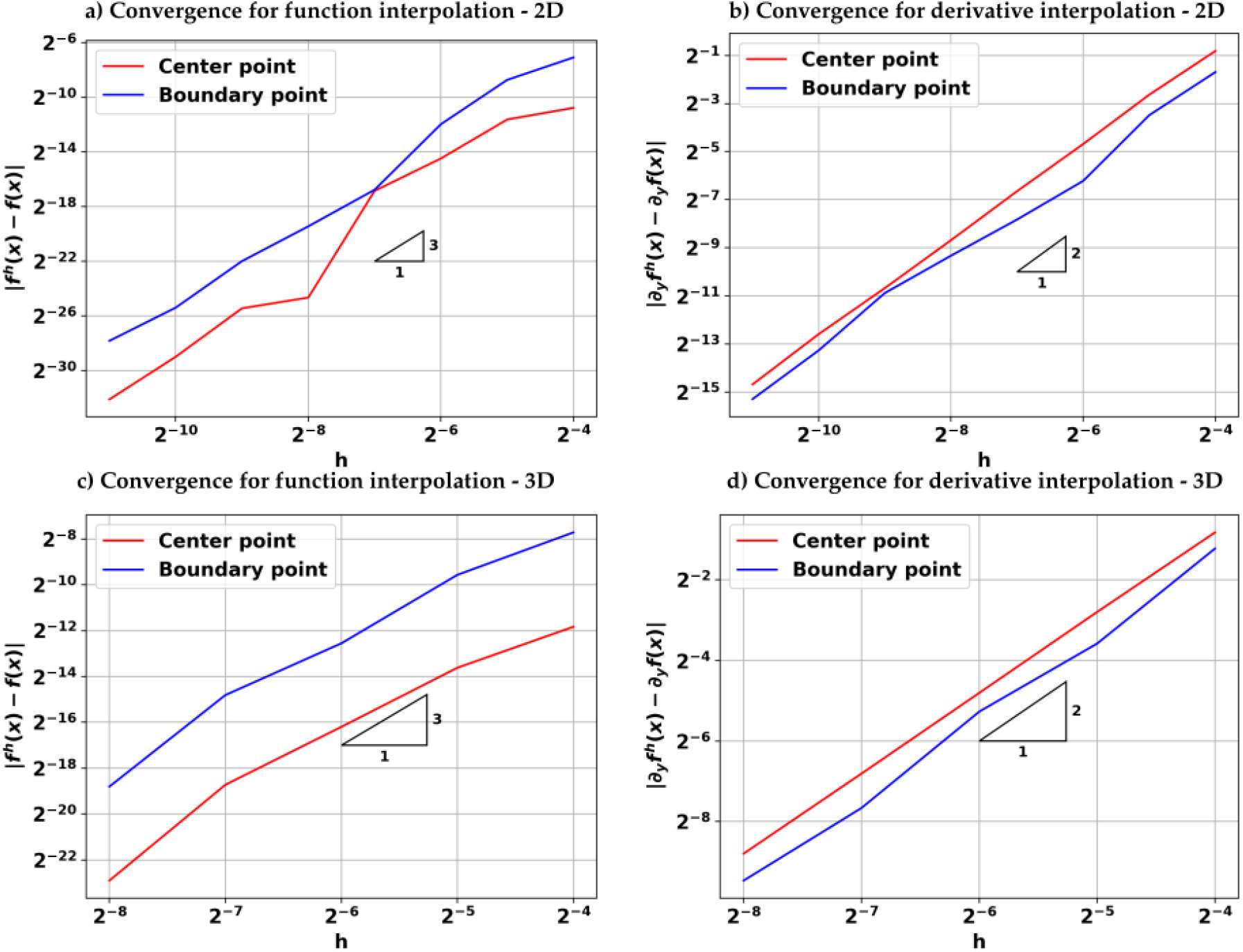
Convergence of the RK interpolation for the center and the boundary point for both 2D and 3D interpolation. Panels a) and b) illustrate the convergence profile for RK interpolation in 2D showing O(h^3^) and O(h^2^) for function and first derivative interpolation respectively. Panels c) and d) illustrate the convergence profile for RK interpolation in 3D illustrating the same convergence rate as the 2D counterpart.

### 3.2 Comparing flow-induced force estimates for immersed and body-conforming meshes

Accurate computation of flow-induced forces on the immersed particles is essential for correctly predicting particle trajectories, such as embolus trajectories for our application of interest in this study. Here, we validate the proposed fluid-solid interaction method for a non-spherical immersed particle, by comparing the predicted flow-induced forces from our immersed formulation against those obtained using a typical body-fitted conforming discretization. Specifically, we consider a solid particle held fixed at an arbitrary orientation to the flow, which helps us achieve a detailed quantitative comparison against a body-fitted discretization without having to re-mesh the fluid domain around the particle. For applications of interest such as that of embolus-hemodynamics interactions, this formulation can be interpreted as resolving the interactions in the embolus’s frame of reference. The model problem setup is illustrated in Figure 4. We specifically compute the fluid forces, by integrating stresses around the tilted ellipsoidal particle within a cylindrical vessel. The ellipsoidal geometry has minor axes radius *r*_*a*_ = 0.05 length units; and major axis radius *r*_*b*_ = 0.075 length units. The embolus is placed 0.2 length units away from the inlet (*along the z direction*), and 0.2 length units away from the bottom wall (*along the y direction*), and symmetrically placed in between the side walls (*x* direction); as indicated in Figure 4. Finally, the ellipsoidal particle is angled such that the major axis is 45^*?*^ to the vessel center-line. The fluid density is set to *ρ* = 1 and the dynamic viscosity is set to *µ* = 10^−3^. We impose a pulsatile inlet flow with a parabolic flow profile defined as follows:

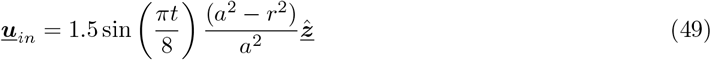

where *a* = 0.205 being the cylindrical vessel radius, and 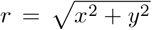 being the radial distance on the inlet face from the inlet center. This formulation gives the Reynolds number with respect to the embolus at *O*(10^2^) at the peak of the pulsatile flow. We define drag force along the *z*-direction, and an effective lift force along the *y*-direction. In the body-conforming discretization case, the local mesh size around the ellipsoidal particle is set to *h* = *r*_*a*_*/*20. For the immersed finite element case with RK interpolation, we progressively refine the mesh until it matches the mesh resolution of the mesh-conforming case. The coefficient of drag *C*_*D*_ in this case is defined as:

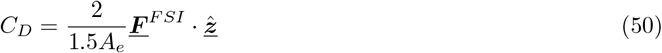

and coefficient of lift *C*_*L*_ defined as:

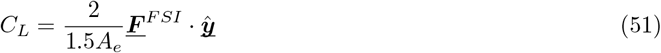

where *A*_*e*_ is the approximate projected area of the ellipsoid defined as *A*_*e*_ = *πr*_*a*_*r*_*b*_; and ***<u>F</u>*** _*FSI*_ represents the computed force from the fluid obtained by integration of stresses along the ellipsoid for both immersed and body-confirming discretizations. Figure 5 a1 and b1, respectively, illustrate the coefficients of lift and drag computed from our proposed immersed method, clearly demonstrating its convergence toward the corresponding coefficient values as obtained from the body-conforming discretization case. This is further substantiated in Figure 5.a2. and 5.b2., which show that the predicted flow-induced force coefficients converge linearly towards those obtained from the more resolved mesh-conforming case. This analysis establishes a numerical basis for assessing the ability of our immersed RK interpolation formulation to reproduce the flow-induced forces on an immersed object.

**Figure 4:**
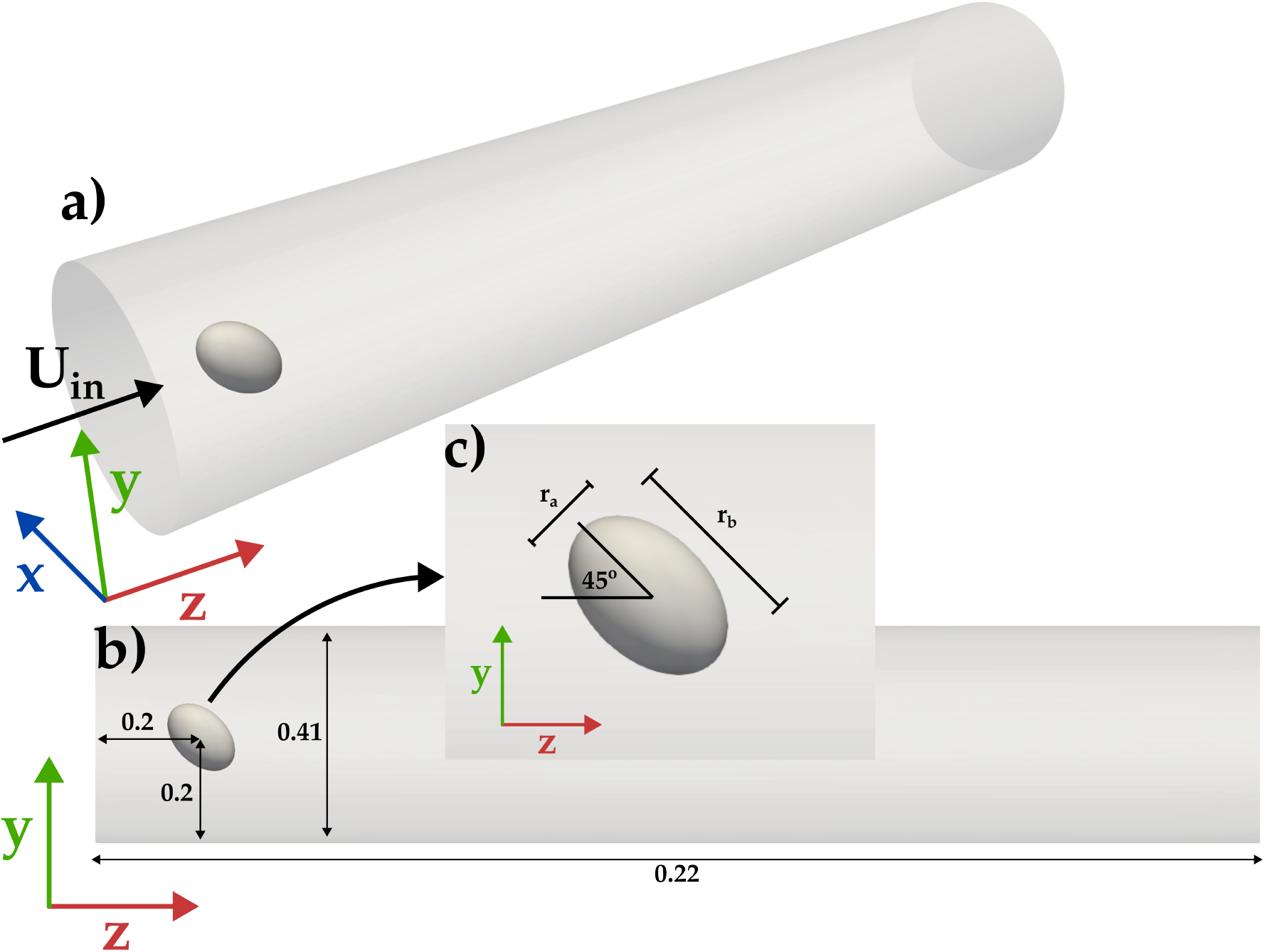
Schematic illustrating the validation problem case of an immersed ellipsoidal embolus inside a cylindrical vessel.

**Figure 5:**
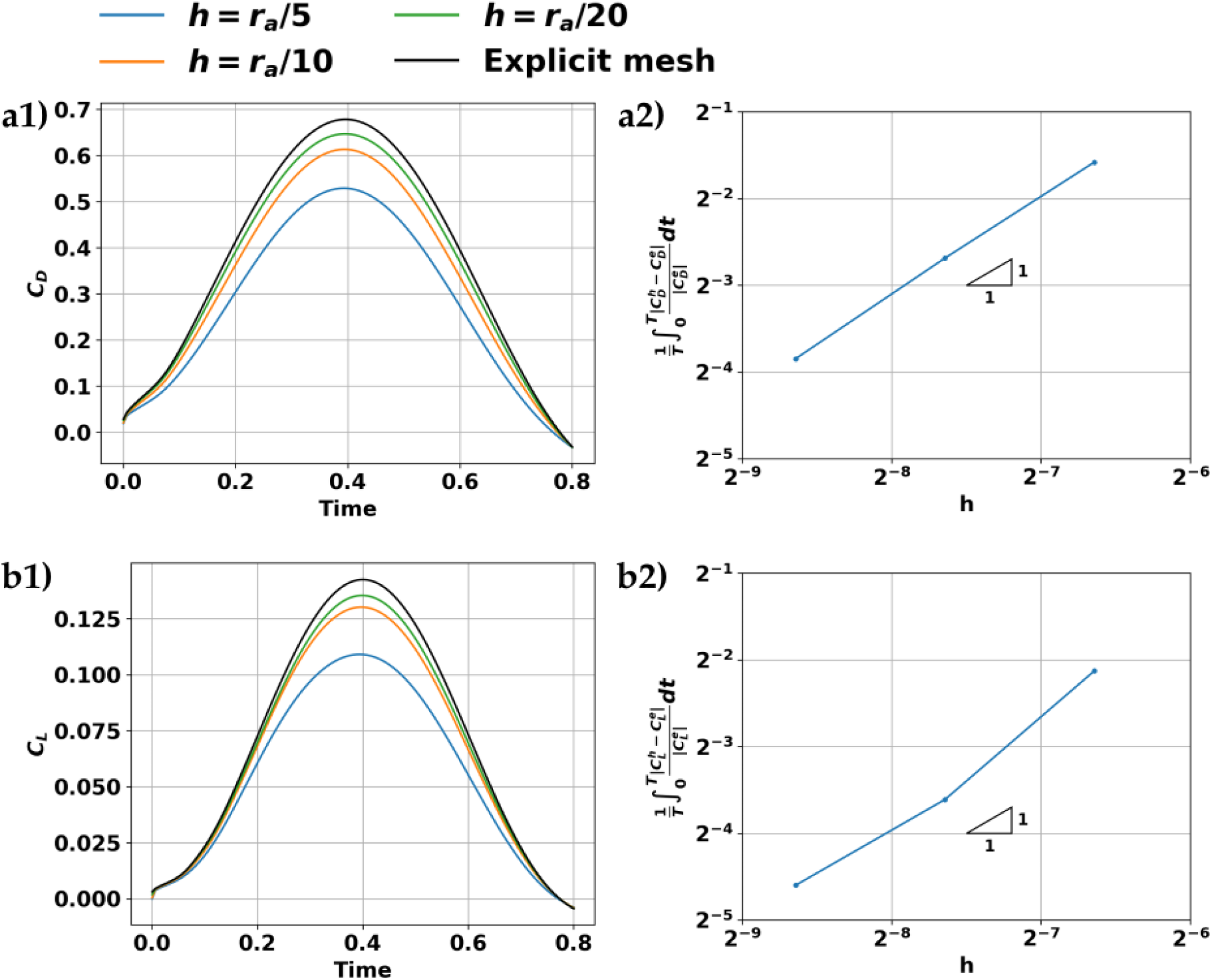
Convergence behavior of the predicted hemodynamic forces on a stationary ellipsoidal embolus. Panels a1 and a2 illustrate the convergence behavior of the coefficient of drag showing linear convergence with local mesh refinement; Panels b1 and b2 also illustrate the convergence behavior of the coefficient of lift showing linear convergence with local mesh refinement.

**Figure 6:**
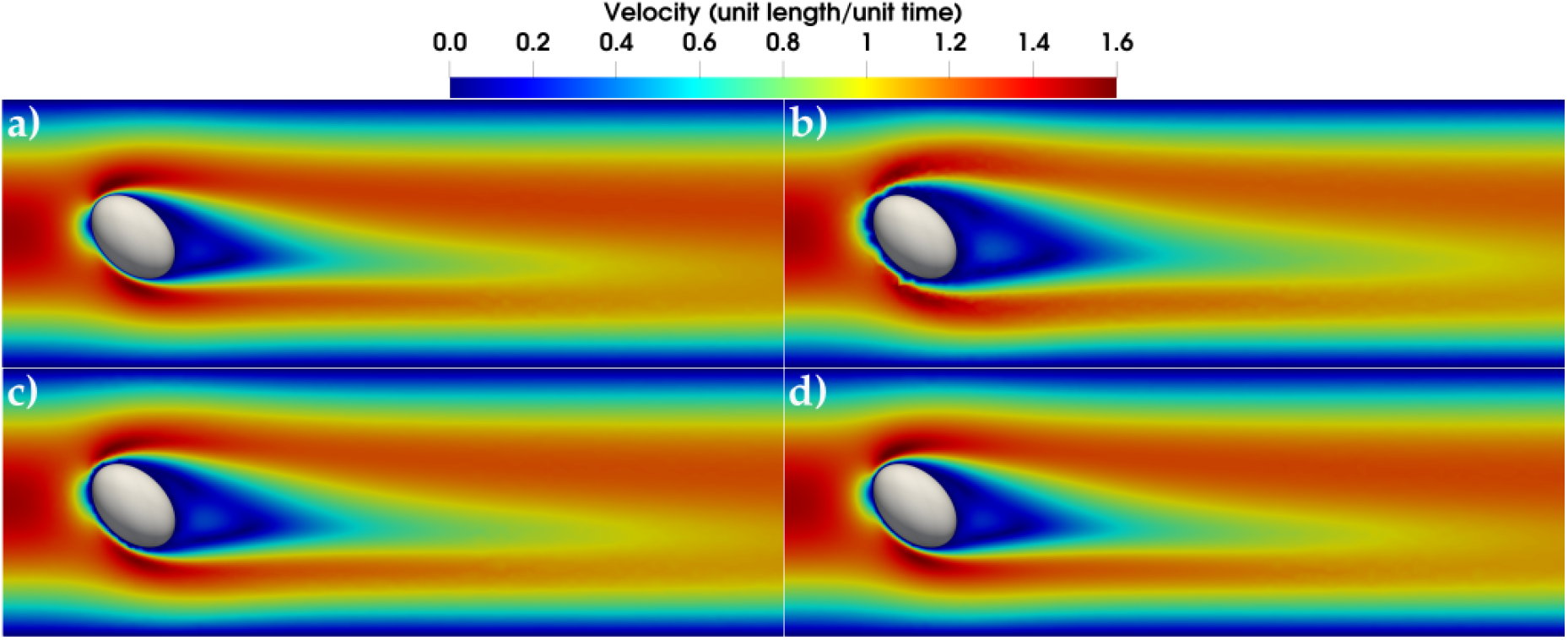
Velocity field of the sample problem along the vessel axis. Panel a) illustrates the explicitly mesh ellipsoidal embolus, and panels b), c), and d) show the velocity field with successively refined mesh sizes h = r_a_/5, h = r_a_/10, and h = r_a_/20 respectively showing that the artificial boundary layer thickness converge to the explicitly mesh along the mesh refinement process.

### 3.3 Benchmark case of a rigid particle falling in a confined 3D channel

We present further numerical demonstration and validation of our immersed finite element framework using a benchmark scenario involving a rigid spherical ball falling in a viscous fluid contained in a confined container. Specifically, we simulate the case representing the experimental study involving a steel ball of radius 1.5 mm falling in silicone oil as reported in [48]. The spherical steel ball has material density of *ρ*_*p*_ = 7.60 g/cm^3^, and the ball is tracked as it falls within a cuboidal container with dimensions 1.0 *×* 1.0 *×* 5.0 cm. The container is filled with silicone oil, assumed to be an incompressible fluid, with kinematic viscosity *ν* = 1000 cSt, and density of *ρ* = 0.97 g/cm^3^. For this study, we did not simulate the particle trajectory as it collide with the container bottom wall, and post-collision. This is because an accurate simulation of the particle-wall interaction requires resolving the particle-wall contact forces with measured restitution coefficients in presence of lubrication effects, which was not the objective of this analysis. Instead, here we focus on the motion of the ball prior to collision with the bottom wall, and ensuring the ball reached terminal velocity. With the material properties considered here, the spherical ball’s equivalent momentum response time is estimated at 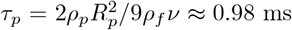,and the terminal velocity is reached at less than 1.0 sec of physical time. For quantitative verification of our methodology, we compared the predicted terminal velocity of the steel ball against those from experimental measurements. Figure 7 presents the results from this verification exercise. Figure 7.a. illustrates the error in the predicted terminal velocity (*compared to experimental values*) with respect to successive mesh refinements - indicating that the terminal velocity predictions converge linearly to the experimental values ***v***_*exp*_ as the mesh size *h* is refined. This is commensurate with the observation that the immersed RK interpolation method for computing the flow-induced forces are observed to also converge linearly with *h* as outlined in Section 3.2. The position of the ball, and the flow velocity around it, at 4 successive instants in time starting from *t* = 0 (*initial instant when the ball starts dropping*), is illustrated in Figure 7.b1-b4. The computed velocity patterns reflect the reported patterns in the original experimental study [48]; as well as flow physics expectations of a low Reynolds number (*Re*≈ 0.03) and low Stokes number flow around a sphere.

**Figure 7:**
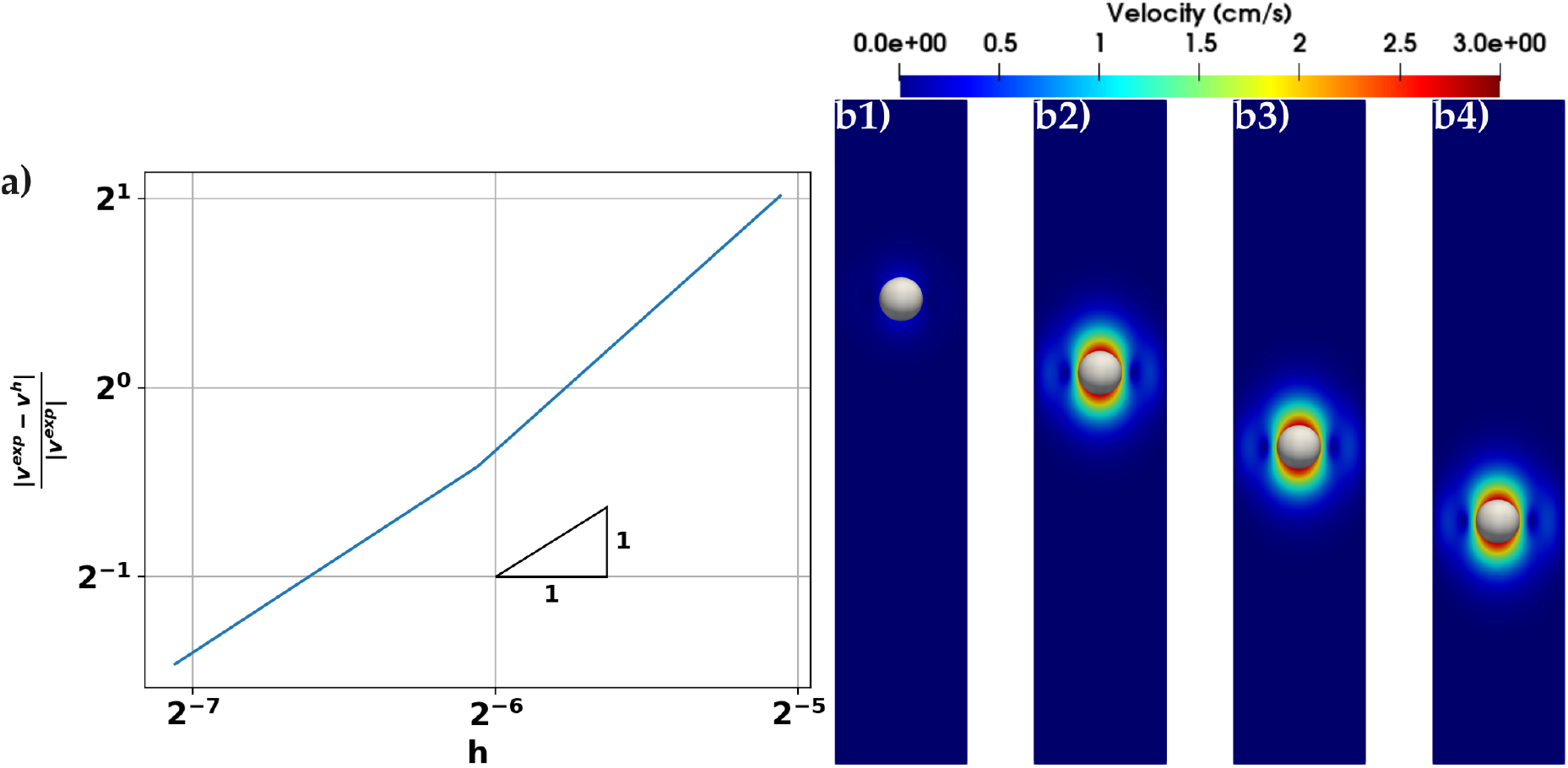
Resulting simulation of the spherical particle falling in a channel. Panel a) show the convergence rate of the spherical particle terminal velocity compared to experimental results in [48] showing the predicted velocity linearly converging to experimental data. Panels b1) to b4) depict the velocity field at time instances 0s, 0.2s, 0.4s and 0.6s respectively depicting the particle reaches terminal velocity very quickly consistent with results shown in [48]

## 4 An in silico study of large vessel occlusions

### 4.1 Description of in silico study setup

We used the computational framework to conduct a parametric numerical study on embolus-hemodynamics interactions for large vessel occlusion scenarios. Specifically, we conducted simulations of a single large embolus (*that is, embolus diameter to vessel diameter ratio greater than 50%*) navigating physiologic pulsatile blood flow across two models of the human carotid bifurcation. The human carotid bifurcation was chosen because of its critical role as a pathway segment for extra-cranial embolic fragments to enter the supplying vessels into the Circle of Willis in the brain (*subsequently leading to a stroke, or related cerebrovascular occlusive event*). The first of these comprised an idealized Y-bufircation model with vessel diameters representative of human carotid vessels. The second model comprised an anatomically realistic carotid bifurcation model derived using vessel segmentation from a contrast-enhanced CT angiography image. This image was obtained from an existing in house database of CT images with de-identified patient records, and was used only for secondary retrospective computational analysis. Hence, no additional IRB documentation was required. For both models, blood was assumed to be a Newtonian fluid with dynamic viscosity *µ* = 4.0 cP and bulk density *ρ* = 1.06 g/cc. A physiological pulsatile flow profile based on measured flow data from [49] was specified at the inlet into the common carotid artery for both; with the flow profile spatially mapped on to the inlet face in form of a parabolic shape. The outflow boundary conditions are set as free-outflow, often referred to as ”do nothing” boundary condition at both outlets. The simulations were allowed to run until the embolus reaches half-way point of either of the two branching vessels (*external and internal carotid arteries*); or until the embolus effectively occludes fully one of the vessels (*tracked by identifying how the embolus contacts the vessel walls*). For a parametric study, we considered two different sizes of ellipsoidal emboli for the anatomical carotid bifurcation geometry, denoted as *E*1 and *E*2; and another two corresponding spherical emboli of equivalent volume: *S*1, and *S*2. The major and minor axes dimensions of each of these cases are compiled in Table 1. Additionally, for the idealized Y-bifurcation geometry, we considered the embolus sizes corresponding to *E*1, *E*2, and *S*1; but eliminated spherical embolus of size *S*2 since the idealized common carotid artery diameter is incidentally smaller than this size, leading to a full occlusion and an ill-posed simulation case. This leads to a sum total of 7 different embolus-hemodynamics interaction models. Vascular geometries for each model were discretized into linear tetrahedral elements with an average mesh size of 0.34 mm for idealized model, and 0.45 mm for anatomical model. Each simulation comprised numerical integration with a time-step size of 10^−3^ sec. For each simulation, the ellipsoidal embolus was initially oriented such that its major axis is normal to the vessel inlet face.

**Table 1:** Table showing the emboli major axis dimensions used in this study. Emboli denoted E1 and E2 are ellipsoidal emboli, and the emboli denoted as S1 and S2 and spherical emboli with equivalent volume to the ellipsoidal emboli.

| Embolus | $r_1(mm)$ | $r_2(mm)$ | $r_3(mm)$ |
| --- | --- | --- | --- |
| $E1$ | 1.00 | 1.00 | 2.00 |
| $E2$ | 2.00 | 2.00 | 4.00 |
| $S1$ | 1.26 | 1.26 | 1.26 |
| $S2$ | 2.52 | 2.52 | 2.52 |

### 4.2 Embolus hemodynamics interactions in an idealized bifurcating vessel

Figure 8 and Figure 9 illustrate snapshots of the embolus with background hemodynamics at four successive instants in time for the simulations involving the idealized bifurcation model. In each case, the embolus was initially positioned along the centerline of the common carotid at the inlet, causing the embolus to move straight along the centerline until reaching the bifurcation region. At the final time instances for all cases, the embolus is observed to move into the internal carotid artery segment (*top branch in the bifurcations in Figure 8- 9*). This is mainly due to the bifurcation geometry for this idealized model having inherenst asymmetry, being slightly off-axis relative to the inlet branch (*common carotid*) centerline, to account for realistic vessel diameter ratios between the internal and external carotid branches. This geometry induces a collision force on the embolus at the bifurcation wall, turning and directing it towards the internal carotid branch. We note that, the trends obtained here are consistent with a previous study using one-way coupled models [15] on an idealized carotid bifurcation model, indicating that emboli show preferential pathway towards the large vessel (ICA). Additionally, although the ellipsoidal emboli *E*1 and *E*2 were initially aligned along the common carotid centerline, the emboli are observed to be off-axis relative to the branch vessels. This misalignment is caused due to torque experienced by the embolus during collisional interaction with the vessel wall at the bifurcation, which leads to tumbling dynamics of the embolus in the vessel. Figure 10.b. presents the time-history of the the embolus angular velocity for the emboli modeled within the idealized bifurcation geometry cases. The observations show no rotation until embolus collision with the off-center bifurcating vessel wall, following which the torque from the collisional interactions lead to a spike in angular velocity. This effect is more pronounced in the ellipsoidal emboli due to their longer moment arm at points of embolus-wall interactions. Furthermore, we observe that the larger ellipsoidal embolus (*E*2) exhibits a smaller angular velocity during and post collision, because its minor axis diameter is comparable to the vessel’s diameter. This creates a near-wall lubrication effect that resists tumbling. In contrast, the smaller ellipsoidal embolus (*E*1) boundary is farther from the vessel wall during tumbling, resulting in a greater angular velocity after the collision. Finally, for the case *E*2 in the idealized bifurcation geometry, at the final time-instance, we are able to nearly resolve a full vessel occlusion event in the internal carotid branch, as evidenced in Figure 9.a4. As the embolus occludes the vessel, we see our simulation recovering the initiation of flow re-routing from the occluded vessel to the patent vessel, as indicated by measured flow-rates at the outlets presented in Figure 10.b. In reality the precise quantitative nature of the flow routing may be different based on actual biophysical interactions of vessel wall with embolus.

**Figure 8:**
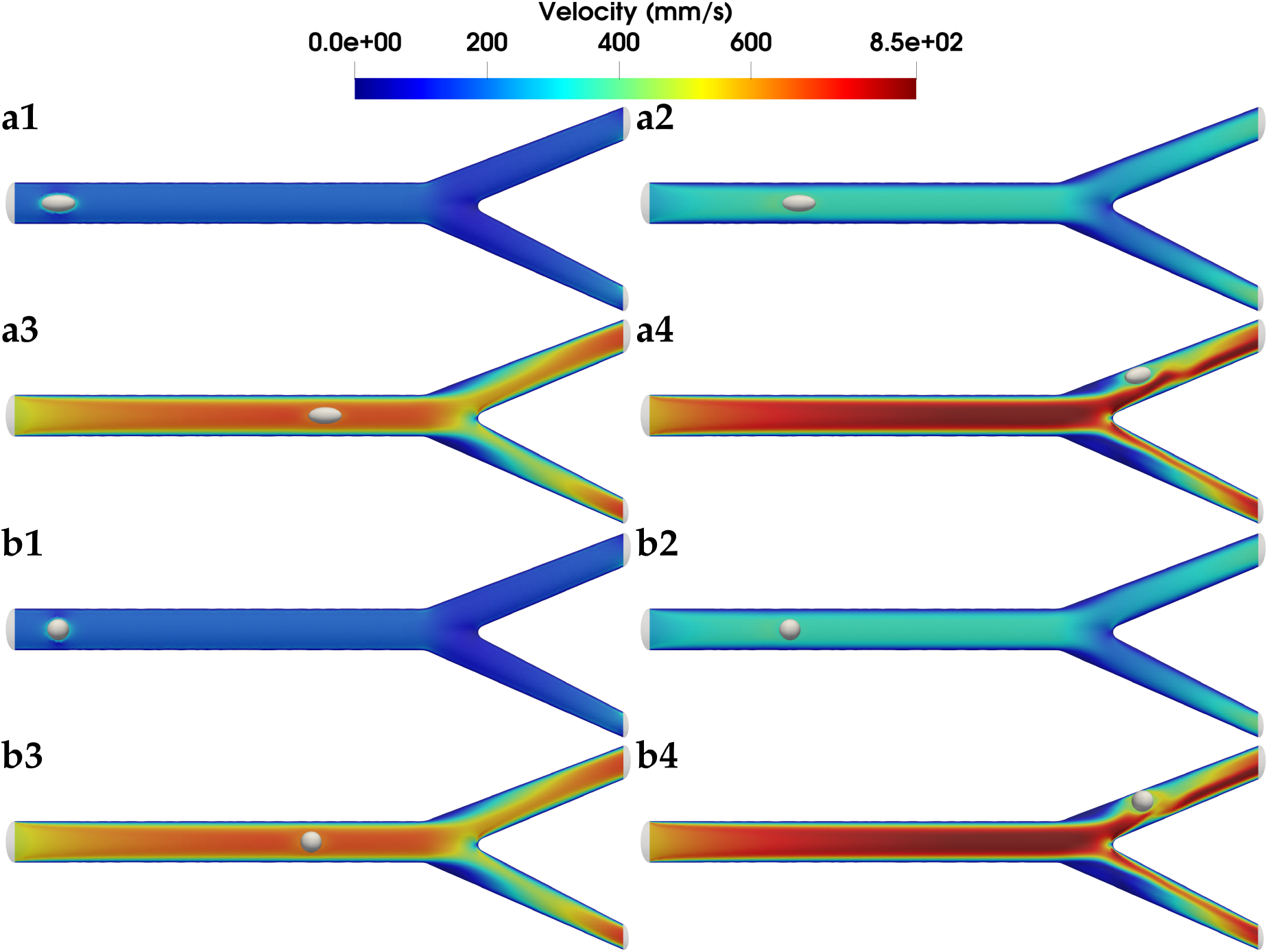
Velocity field at the center plane of the idealized Y-bifurcation for the small spherical embolus case (S1) (shown in Panels a) and the small ellipsoidal embolus case (E1) (shown in Panels b) at four time instances; Panels a/b1 at time t = 0 s, a/b2 at time t = 0.04 s, a/b3 at time t = 0.08 s, and a/b4 at time t = 0.12 s

**Figure 9:**
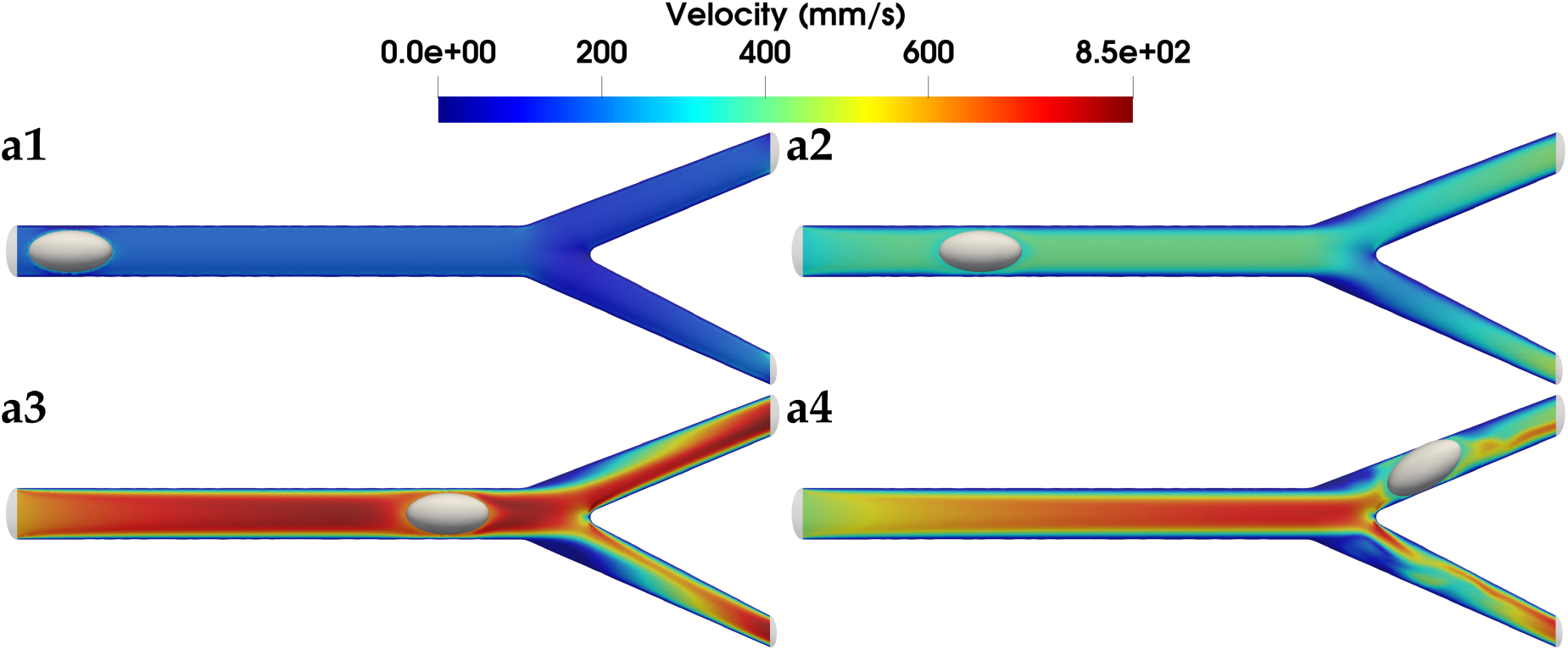
Velocity field at the center plane of the idealized Y-bifurcation for the large ellipsoidal embolus case (E2) at four time instances; a1 at time t = 0 s, a2 at time t = 0.05 s, a3 at time t = 0.10 s, and a4 at time t = 0.15 s

**Figure 10:**
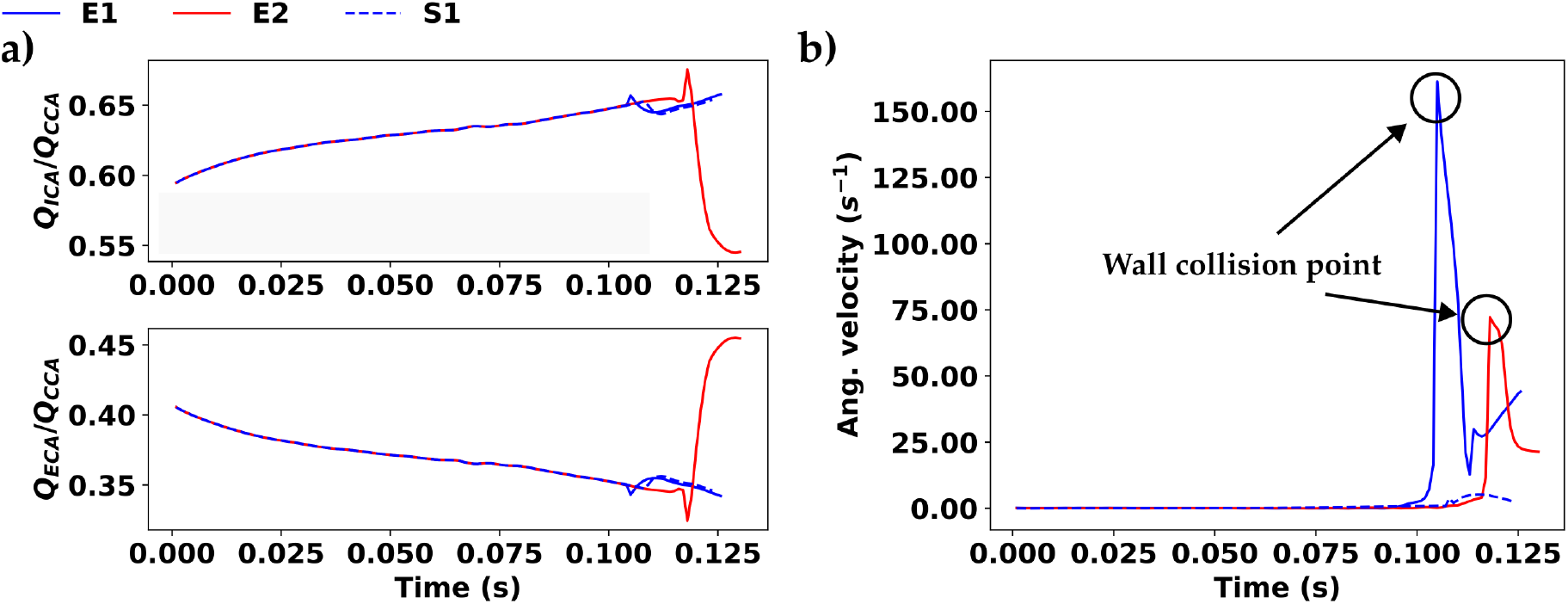
Panel a) depicts the ratio between the outlet volume flow rate leaving each outlets (internal carotid artery (Q_ICA_) and the external carotid artery (Q_ECA_)) with the total inlet flow rate through the common carotid artery (Q_CCA_) in the idealized Y bifurcation case. In the large ellipsoidal embolus (E2) case, the embolus traveled into the ICA and colliding with the vessel wall resulting in the flow re-routing from the ECA to the ICA deviating from other cases. Panel b) illustrate the spike in the embolus’ angular velocity indicating the point of embolus collision with the vessel wall.

### 4.3 Embolus hemodynamics interactions in an anatomical carotid bifurcation

Figure 11 and Figure 12 illustrate the flow streamlines and embolus movement at snapshots across four successive instances in time for the simulations involving anatomical carotid bifurcation geometry. We observe that the interaction of pulsatile flow with curved anatomical vessel geometries leads to highly nonlinear flow patterns, which are locally altered as the embolus traverses the vessels. In all cases, we note embolus movement towards the larger branch vessel (*the internal carotid*) after hitting the bifurcation (*again, similar to findings in prior work such as [15]*). However, compared to the idealized models, the ellipsoidal emboli show significant tumbling motion as they approach the bifurcation point. The embolus-wall interactions and resultant tumbling dynamics is illustrated using time-history of angular velocities presented in Figure 13. We note that both ellipsoidal and spherical emboli exhibit non-zero angular momentum, due to the curvature and tortuosity of the anatomical bifurcation geometry. For instance, as the ellipsoidal embolus collides with the vessel wall, the angular acceleration is high, leading to a spike in angular velocity (*as noted at t* = 0.05 *sec and t* = 0.22 *sec in Figure 13.b*.). Additionally, for the larger embolus *E*2, larger angular momentum prior to collision at the bifurcation wall leads to secondary tumbling, preventing the angular momentum from reducing immediately post collision. We observe that this secondary tumbling causes the large ellipsoidal embolus to orient orthogonally to internal carotid centerline, thereafter leading to a full vessel occlusion scenario as the ellipsoid lodges within the vessel. The lodging effect can be observed by plotting the volume flow rate ratio between the two outlets relative to the inlet flow rate, as shown in Figure 13.a. Before the emboli reach the bifurcation point, the outlet flow rate remains consistent across all embolus shapes and sizes. More flow is directed toward the ICA outlet compared to the ECA due to the ICA’s larger diameter, resulting in less hydraulic resistance to flow. For emboli *S*1, *S*2, and *E*1, the flow rate ratio remains largely consistent throughout the simulation. However, the large ellipsoidal embolus *E*2 tumbles and eventually lodges within the ICA, causing flow to be redirected from the ICA to the ECA. This effect is reflected in a sharp drop in *Q*_*ICA*_ at approximately *t* = 0.33s in Figure 13.a. In summary, these findings demonstrate that our methodology can capture flow resistance and flow rerouting due to the presence of a large occlusive embolus characteristic of LVOs.

**Figure 11:**
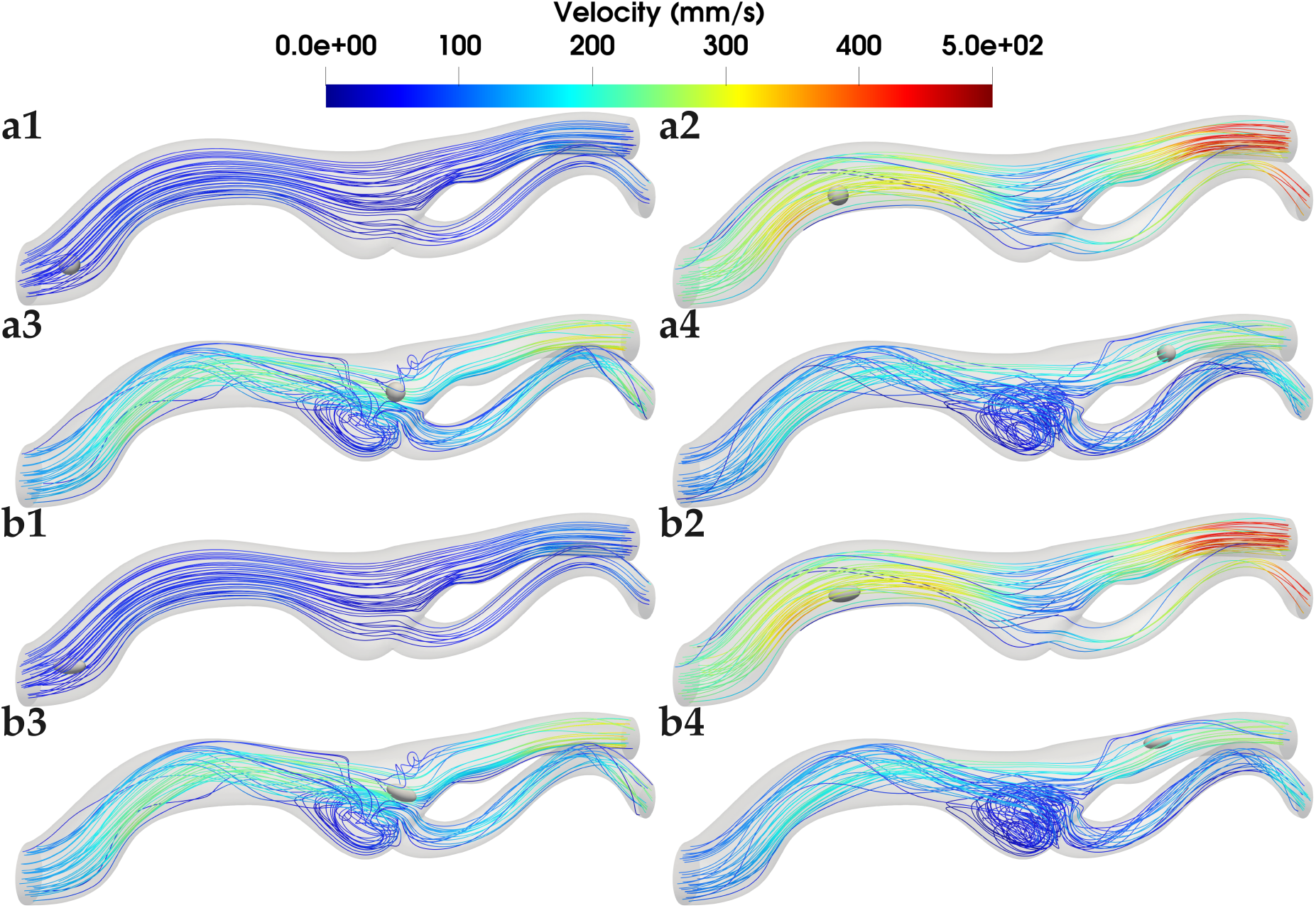
Velocity streamline and embolus position for the small spherical embolus case (S1) shown in panels a1 − a4 and small ellipsoidal embolus case (E1) shown in panels b1 − b4 at four time instances; Time instances are demarcated as panel number a/b1) through a/b4) corresponding to time instance t = 0 s t = 0.1 s, t = 0.2 s, and t = 0.3 s respectively.

**Figure 12:**
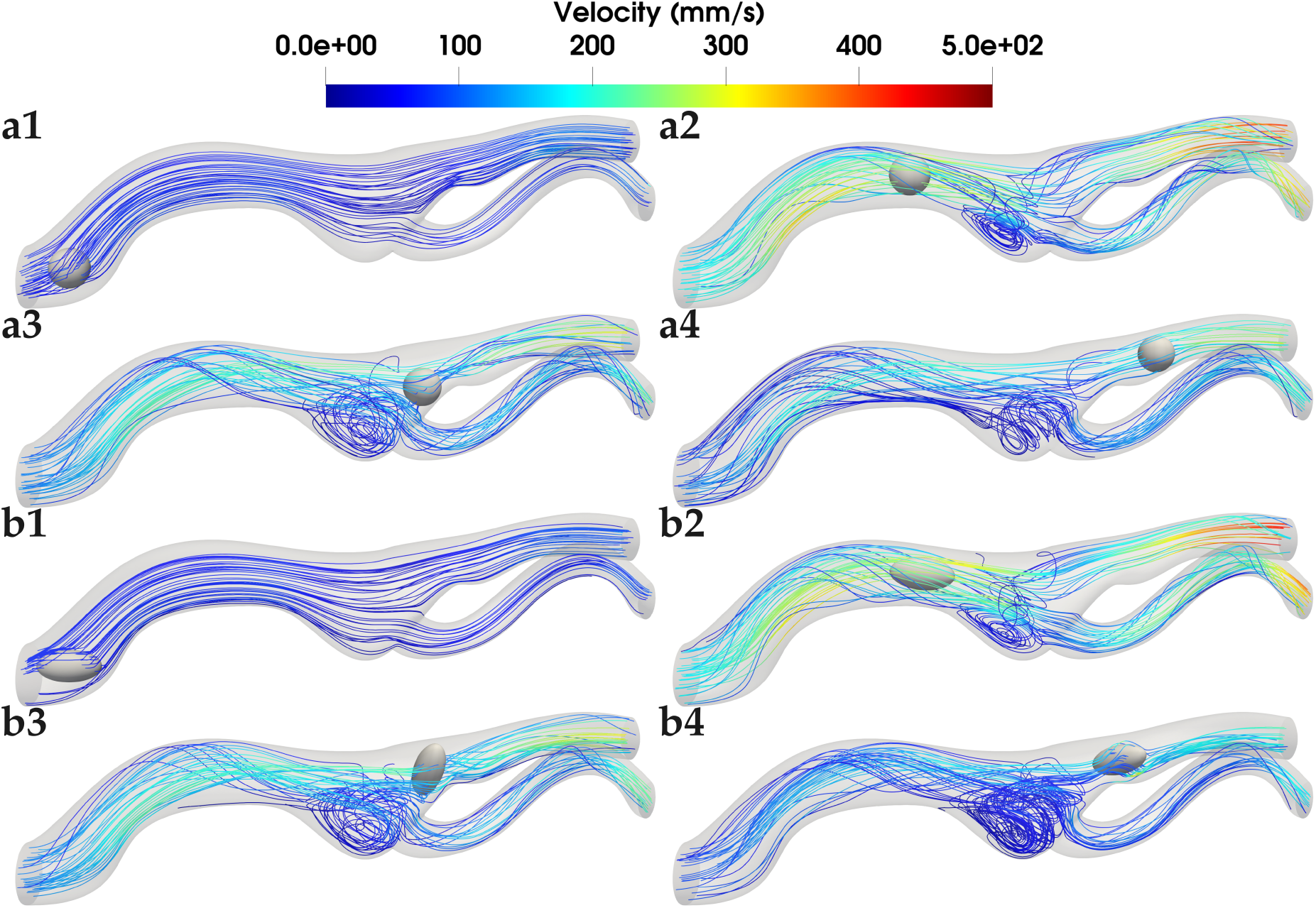
Velocity streamline and embolus position for the large spherical embolus case (S2) shown in panels a1 − a4 and large ellipsoidal embolus case (E2) shown in panels b1− b4 at four time instances; Panels a1 − a4 depict embolus S2 trajectory at time instance t = 0 s, t = 0.14 s, t = 0.28 s, and t = 0.42 s respectively. Panels a1 − a4 depict embolus E2 trajectory at time instance t = 0 s, t = 0.14 s, t = 0.28 s, and t = 0.34 s respectively.

**Figure 13:**
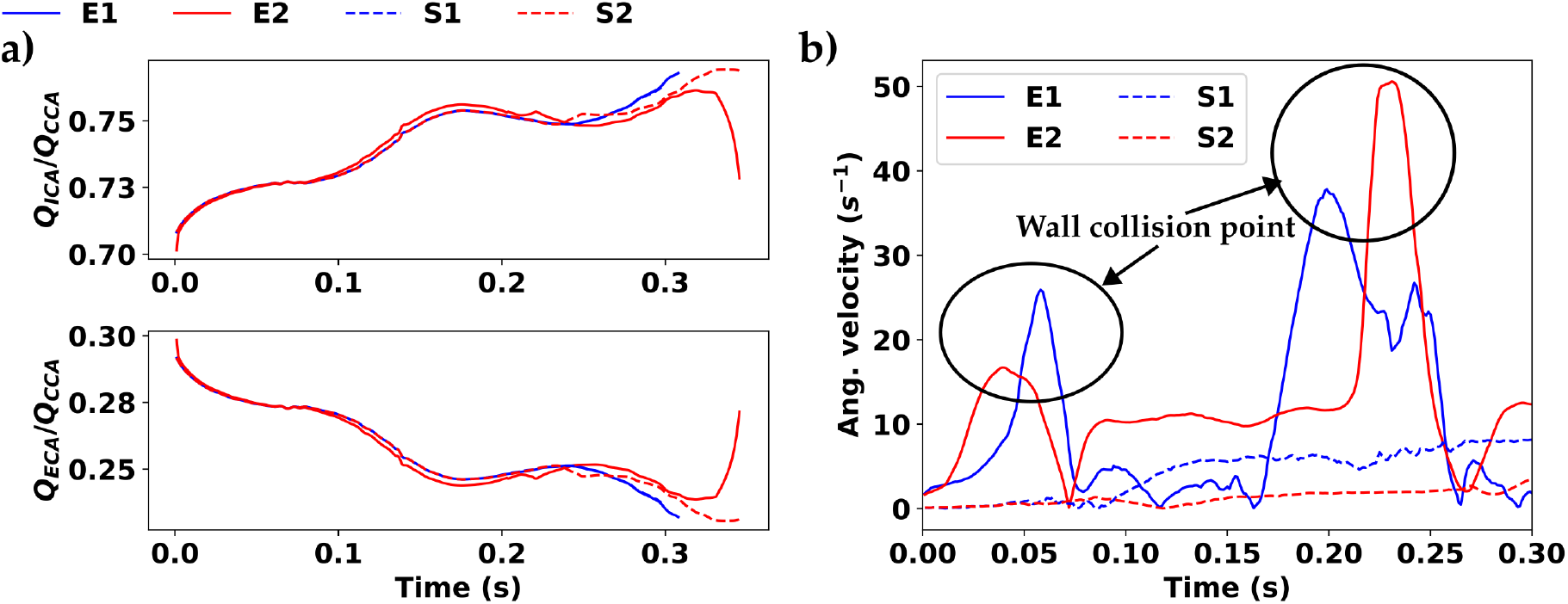
Panel a) depicts the ratio between the outlet volume flow rate leaving each outlets (internal carotid artery (Q_ICA_) and the external carotid artery (Q_ECA_)) with the total inlet flow rate through the common carotid artery (Q_CCA_) in the anatomical bifurcation case. In the large ellipsoidal embolus (E2) case, the embolus traveled into the ICA and colliding with the vessel wall resulting in the flow re-routing from the ECA to the ICA deviating from other cases. Panel b) illustrate the spike in the embolus’ angular velocity indicating the point of embolus collision with the vessel wall first at the initial bend of the anatomical vessel, and second at the bifurcating point.

## 5 Discussion

We presented here a novel algorithmic framework for modeling and simulation of large occlusive emboli in pulsatile hemodynamics within vascular segments. To the best of our knowledge, this is a first-of-its-kind model for non-spherical embolic dynamics relevant for large vessel occlusion (LVO) scenarios. There has been a substantial recent focus on LVO modeling, the majority of which is dedicated to pre-clinical in vivo models [50]. This has been complemented by emergence of image-driven AI/ML models for LVO detection directly based on CT (or equivalent standard-of-care) imaging [51]. None of these leverage any local biomechanical information at the occlusion site, or information on source-to-destination propensity and distribution of emboli. The lack of this information has major consequences on assessment of LVO treatment efficacy using thrombolytics or mechanical thrombectomy, or a combination thereof [52]. As illustrated in the simulations presented here, several complex aspects such as non-linear embolus-hemodynamics interactions, tumbling dynamics of emboli based on wall interactions, local flow routing, lubrication forces, underlie LVO - which are not discernible from imaging and animal models alone owing to challenges of parametric investigations in such modalities. Our framework presented here provides the basis for future in silico LVO models to complement in vivo models.

Furthermore, embolic events often manifest in form of embolic showers, which have implications in determining distal embolization risks during mechanical thrombectomy [23, 53] as well as microembolic infarcts responsible for vascular dementia [6, 54, 55]. While these are not explicitly LVO equivalent cases, the underlying biomechanical event nevertheless involves an embolic occlusion, which can be modelled using the framework developed here. However, the presence of multiple embolic particles necessitates an inter-particle interaction model. For spherical embolic particle, simple particle contact models based on inter-particle penetration and Hertzian elastic contact mechanics will be well suited [38, 39]. However, for scenarios involving non-spherical particles, rapid embolus contact detection is not straightforward, and our SDF-based approach here holds distinct advantages. Specifically, we can project the SDF on the boundary points on each embolus particle relative to every other individual embolic particles. We can employ an accelerated data structure like an Axis-Aligned Bounding Box (AABB) [56] to filter out particles that are too far apart to collide, thereby reducing the actual collision check computations. SDF based collision between objects is in fact commonly employed in computer graphics applications [44, 57–59].

We note that here we show embolus dynamics simulations up to occlusion point. After the embolus gets lodged into the wall, the actual embolus-wall interaction needs to further be augmented using some cohesive interactions, proper computational resolution for which would be a significantly involved task beyond the scope of this study. Additionally, here we have not incorporated the commonly used Windkessel boundary conditions [60] for the vascular flow. In fact, for the kind of scenarios simulated here, the implementation of such outflow boundary conditions need careful attention, because as the embolic particle reaches near-occlusion sizes at a vessel location, the significant level of flow alterations elicit differences from the flow conditions that Windkessel boundary conditions are based on. Furthermore, while our reproducing kernel (RK) implementation provides a robust estimate of flow-induced forces on the particle; for long time dynamics, small errors in forces/accelerations can build up and lead to significant deviations in numerical predictions. This can be mitigated, for example, using numerical correction forces such as those reported in [27].

While our SDF based approach enables efficient contact detection, the actual contact interaction forces will need to account for embolic particle material constitutive model - which is an aspect that we have not explicitly addressed here for conciseness of scope of the study. Thromboemboli are known to exhibit viscoelastic properties [61] therefore, a potential approach is to adopt a viscous Hertzian-type contact model, similar to those described in [38]. If the assumption of negligible deformation is maintained, incorporating this implementation remains straightforward, as the collision force is a simple algebraic expression of collision depth and velocity. The semi-rigid embolus assumption is particularly appropriate for emboli originating from cholesterol plaque rupture as they tend to be stiffer than thromboemboli. However, for softer thromboemboli, significant deformation can occur, which requires a deformable embolus model. Our proposed model can be extended to incorporate deformable emboli using the deformable IFEM framework from [27]. In this case, the 6-DOF formulation presented in Section 2.3 will be replaced with a solid mechanics equation with an appropriate constitutive model for the embolus. The vessel collision detection framework will remain unchanged, as we will continue to use the SDF to accelerate collision detection on the embolus now deformed boundary points. Equation 45 can be seen be seen as a traction boundary condition in the deforming embolus equation.

Finally, the proposed framework utilized a stabilized finite element formulation for hemodynamics [28] which works well for low to moderate Reynolds number regime. Here, we focus on smaller arterial vessels, such as the carotid bifurcation. However, in larger arteries such as the aorta, blood flow can transition to turbulence [62]. In such cases, capturing the smallest flow scales requires either a very fine mesh resolution which is not always computationally tractable, or an appropriate sub-resolution modeling to account for unresolved turbulent effects. In this case, we can modify equations (11) to account for sub-grid flow effect using methods such as the Variational Multiscale (VMS) [63, 64]. While the effect of small scale flow features is captured in the flow at large, the implication of VMS on the IFEM coupling framework remains unclear. However, for the specific application area of embolisms as discussed here, this is not problematic because during occlusion events characteristic to LVOs, flow will be diverted due to additional flow resistance which maintains a low Reynolds number near the occlusion site.

Beyond the focus application of embolus-hemodynamics interactions, we note that the overall computational framework as developed here is fully generalizable to any two-way coupled fluid-particle interaction problem where the particle shape is non-spherical and the background internal flow is occurring within a complex geometry. In this framework, we leveraged a projection of the SDF onto the particle boundary, and used that to generate a distributed contact loading for resolving wall contact with the artery walls. This technical detail is of high significance when considering geometric complexities that commonly arise in real-world biomedical (as well as industrial problems). The inherent superquadric based parametric shape representation enables major advantages in resolving the particle-fluid (in this case, embolus-hemodynamics) interaction mechanisms, while retaining an immersed non-conforming discretization. We note that the standard superquadric shapes can be further modified using Bezier curves to represent the latitudinal and longitudinal exponent functions, thereby providing a high degree of flexibility in representation of arbitrary shapes (referred to as *extended superquadrics*) [30, 65].

## 6 Concluding remarks

In this study, we present a novel algorithm for two-way coupled immersed fluid particle interactions, with applications to large embolus hemodynamics interactions within complex arterial geometries. The two-way interactions are modeled using an immersed finite element method framework, adapted for immersed rigid solids. A signed distance field is employed to detect and compute embolus-vessel wall contact forces, using a paremetric shape representation for the embolus. The two-way coupling enables the simulation of large emboli interacting and potentially lodging within vessels, which is a key characteristic of Large Vessel Occlusion (LVO) stroke. We verified that our proposed model showed good numerical performance and convergence for comparisons against simplified benchmarks, experimental data, and was successfully able to replicate embolus dynamics in anatomical and idealized vascular bifurcation models. To the best of our knowledge this is the first model to capture LVO stroke scenarios and underlying near-occlusive embolus dynamics, although the methodology is generalizable to a wide-range of other fluid-particle interaction applications.

## Supporting information

Anatomical-Embolus-Simulations-LVO-Ellipsoidal

Anatomical-Embolus-Simulations-LVO-Spherical

## Ethics statement

This study used patient data available in house in investigator DM’s research group at CU Boulder in an anonymized form, solely for retrospective secondary computational analysis. No additional Institutional Review Board (IRB) considerations were required.

## Acknowledgements

This work was partially supported by a University of Colorado Boulder Research and Innovation Office Seed Grant award, as well as partially supported by a National Institutes of Health award (R21EB02973) - both awarded to DM. This work utilized the Alpine high performance computing resource at the University of Colorado Boulder. Alpine is jointly funded by the University of Colorado Boulder, the University of Colorado Anschutz, and Colorado State University and with support from NSF grants OAC-2201538 and OAC-2322260. Data storage supported by the University of Colorado Boulder ‘PetaLibrary’ (allocation to DM).

## Author contributions

CT and DM devised the underlying numerical methodology. AK devised incorporation of SDF based algorithms into the methodology. CT led the computational implementation of the methodology, and conducted computational simulations and data analysis. CT and DM prepared the manuscript, including draft review, and editing. AK worked on manuscript review and editing. All authors agreed to the final version of the manuscript as submitted.

## Data, materials, and code disclosure

The complete computational implementation, along with demo case file for the bifurcation models, will be made available as part of the open source finite element flow and transport toolkit named FLATiron Toolkit developed and maintained by DM’s research group at the University of Colorado Boulder. The toolkit is available open source on GitHub at the following link: https://github.com/flowlabcu/FLATiron.

## Supplementary Material Information

### S1 Description of the overall simulation algorithm

#### Algorithm 1

Fictitious domain IFEM

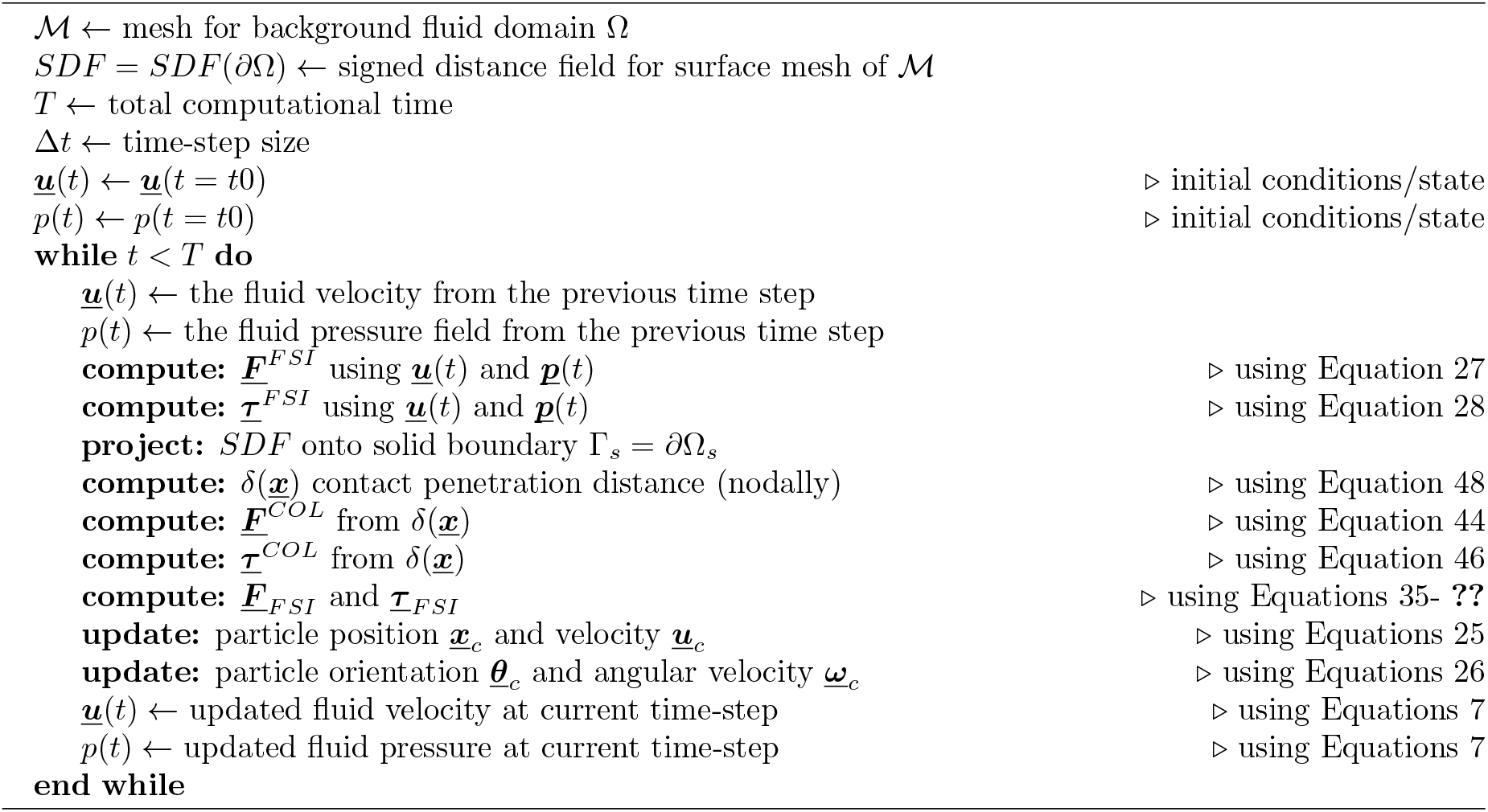

### S2 Numerical details of Gear’s algorithm

Gear’s algorithm consists of two steps: first, for a known position ***<u>x</u>*** and the corresponding time derivatives 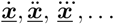 at time *t*, the predictor step estimate the particle’s position its time derivative at time *t* + Δ*t* based on Taylor expansion. The degree of truncation depends on the order of integration algorithm. In this work, we employ the fifth order integration algorithm typically employed in DEM and molecular dynamics [38] defined as:

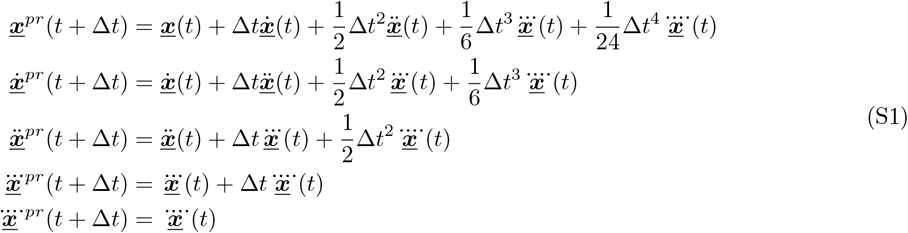

The acceleration due to the applied force and torque terms are computed at *t*+Δ*t* at the start of integration. The computed accelerations at time *t*+Δ*t*, however, usually differ from the predicted value 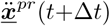.Gear’s algorithm uses the difference between the predicted and corrected acceleration to compute the correction of the velocity and position. Hence, the second step of Gears integration is the correction step as follows: let the difference between the predicted and corrected acceleration be 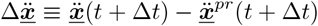.The corrections for the position and the corresponding time derivatives are computed as follows

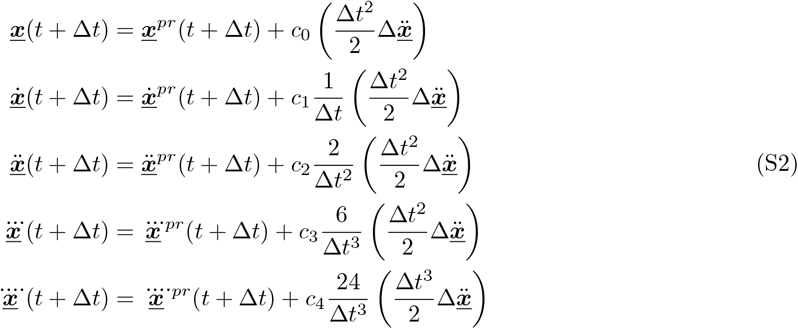

With the coefficients of fifth order Gears integration are as follows

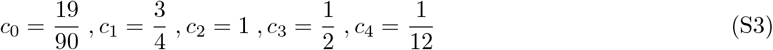

